# CRISPR-Decryptr reveals cis-regulatory elements from noncoding perturbation screens

**DOI:** 10.1101/2020.08.13.247007

**Authors:** Anders Rasmussen, Tarmo Äijö, Mariano Ignacio Gabitto, Nicholas Carriero, Neville Sanjana, Jane Skok, Richard Bonneau

## Abstract

Clustered Regularly Interspace Short Palindromic Repeats (CRISPR)-Cas9 genome editing methods provide the tools necessary to examine phenotypic impacts of targeted perturbations in high-throughput screens. While these technologies have the potential to reveal functional elements with direct therapeutic applications, statistical techniques to analyze noncoding screen data remain limited. We present CRISPR-Decryptr, a computational tool for the analysis of CRISPR noncoding screens. Our method leverages experimental design: accounting for multiple conditions, controls, and replicates to infer the regulatory landscape of noncoding genomic regions. We validate our method on a variety of mutagenesis, CRISPR activation, and CRISPR interference screens, extracting new insights from previously published data.

## Main

Information garnered from pooled CRISPR perturbation screens impacts decisions that have therapeutic implications. Genome-wide knockout and noncoding screens have been used to identify new therapeutic targets, to reveal genes responsible for anti-cancer drug resistance, and to map functional elements in leukemia cell lines.^1,2,3,4^ As researchers in academia and industry make greater use of improving gene editing technologies, computational approaches that tackle the unique challenges posed by their experimental design are of increasing importance. Methods employed for knockout screens are designed to assess the impact of perturbing a genome-wide set of pre-delineated coding regions. ^5,6^ However, analysis of CRISPR noncoding screens, which employ saturated guide libraries to reveal *cis*-regulatory elements, necessitate distinct experimental considerations. Most importantly, classification of functional elements without *a priori* knowledge of their location or size requires integrating information across perturbations within genomic proximity, an aspect that renders existing knockout methods inapplicable to these experimental designs. Literature on methods for analyzing noncoding screens is scarce, with only a single method published that addresses one of the many aspects of noncoding screen analysis.^7^

CRISPR-Decryptr utilizes techniques from Bayesian inference, signal processing, and latent variable models to integrate data and experimental design, allowing the end-user to make precise conclusions about their noncoding screen results (**Figure 1**; *Methods*). A Bayesian hierarchical generalized linear model (GLM) serves as the mathematical formulation from which perturbation-specific effect on phenotype are inferred^8, 9^. The model leverages experimental conditions, controls, and replicates in a single numerical procedure implemented with Markov Chain Monte Carlo, allowing for rigorous statistical treatment of parameter uncertainty (*Methods 2.2*). Effects are mapped to a base-by-base level of granularity through a Gaussian process-based model (*Methods 2.4*)^10^. This deconvolution fully accounts for guide-specificity, off-target effects and, if applicable, double-strand break (DSB) repair uncertainty (*Methods 2.3*)^11, 12^. A hidden semi-Markov model (HsMM) incorporates spatial information to decode the latent regulatory landscape of interest, revealing enhancers and silencers in the noncoding genome (*Methods 2.5*)^13^. Regulatory element calls and guide-specific effects are exported in bioinformatics file formats such as Browser Extendable Data (.bed) and Wiggle (.wig) that can easily be explored in genomic visualization software such as the Integrative Genomics Viewer (IGV)^14^.

**Figure 1:**
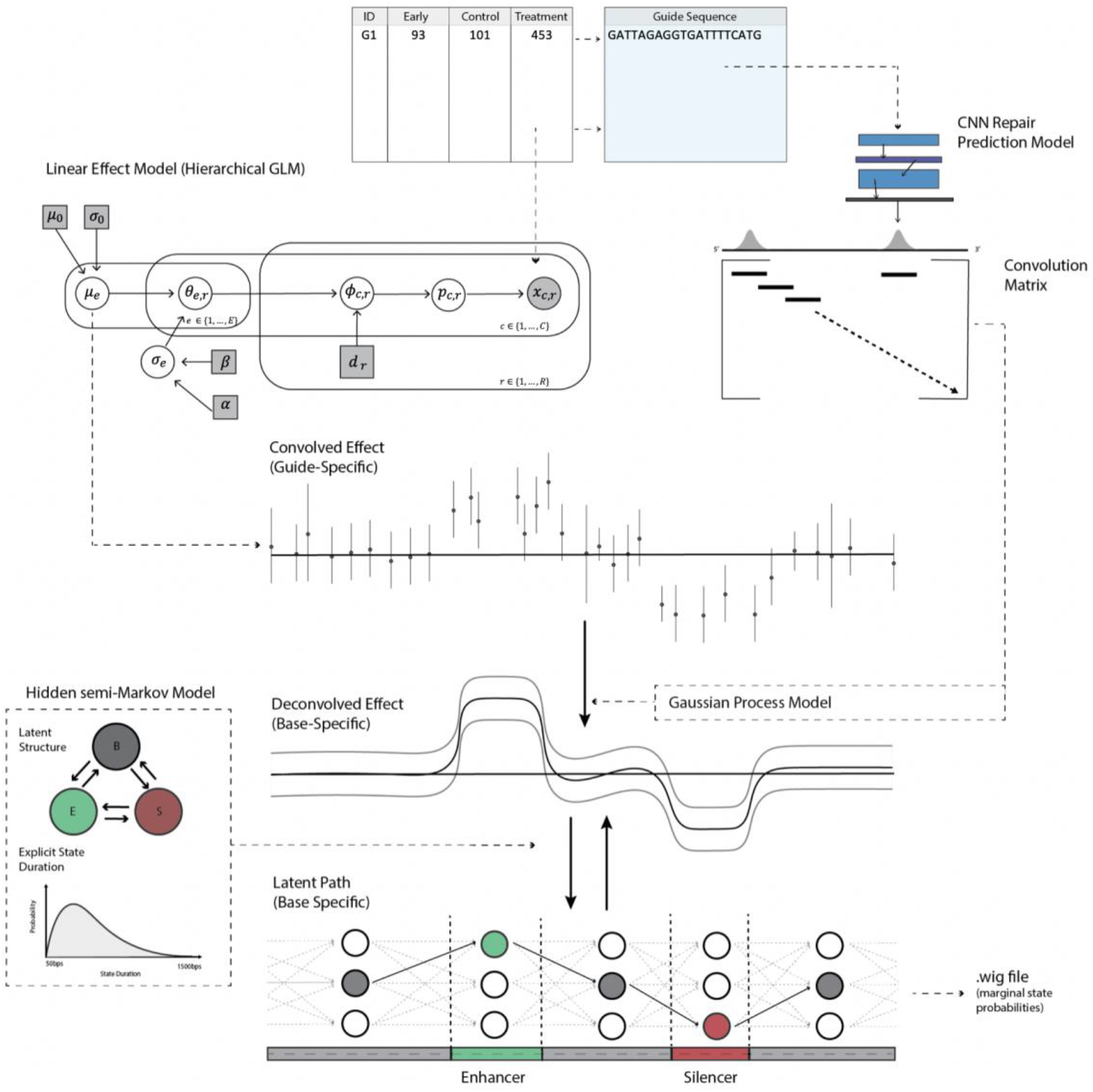
Overview of the CRISPR-Decryptr method for the analysis of noncoding screens. The hierarchical GLM infers pertubation-specific regulatory effect on phenotype from raw guide RNA (gRNA) counts (*top left*). Guide RNA sequences are used to construct a convolution matrix accounting for specificity, off-target effects, and repair uncertainty in the case of mutagenesis screens (*top right*). Finally, iterating between Gaussian Process deconvolution and HsMM training and prediction reveals base-specific effects and ultimately the latent state path of interest (*bottom half*).

We validated CRISPR-Decryptr on noncoding screens of distinct experimental designs, including CRISPR mutagenesis, CRISPR activation (CRISPRa), and CRISPR interference (CRISPRi) screens (**Figure 2**) ^4, 15, 16^. In the CRISPR mutagenesis screen (Canver *et al.*), three intronic DNAse hypersensitivity sites (DHS) within *BCL11A* were perturbed in human umbilical cord blood-derived erythroid progenitor (HUDEP) cells^15^. These sites, termed DHS +62, +58, and +55, are known to impact fetal hemoglobin (HgF) levels from prior published research, with the enhancer identified in DHS +58 having proven a successful therapeutic target in two patients with hemoglobinopathies.^17^ When applied to this dataset, CRISPR-Decryptr produced regulatory state calls in agreement with the original analysis (**Figure 2A** and **Supplementary Figure 3.1.1**). The CRISPR activation screen we re-analyzed (Simionov *et al.*) targeted the *IL2RA* and *CD69* gene loci in Jurkat T-cells.^16^ To measure phenotypic change, the FACS sort cells into a “negative”, “low”, “medium”, and “high” bins of IL2RA and CD69 based on expression levels. Analysis of the two gene loci with CRISPR-Decryptr recalls the enhancers from the original analysis, as well as novel putative enhancers that are correlated with DNAse-seq and H3K27ac from the Jurkat-T Cell line (**Supplementary Figures 3.2.2** and **3.2.3**). Finally, the re-analysis of the Fulco et al. CRISPRi screen of the GATA1 gene loci revealed similar regulatory element calls to the original analysis.^4^ (**Figure 2C** and **Supplementary Figure 4.3.1**).

**Figure 2:**
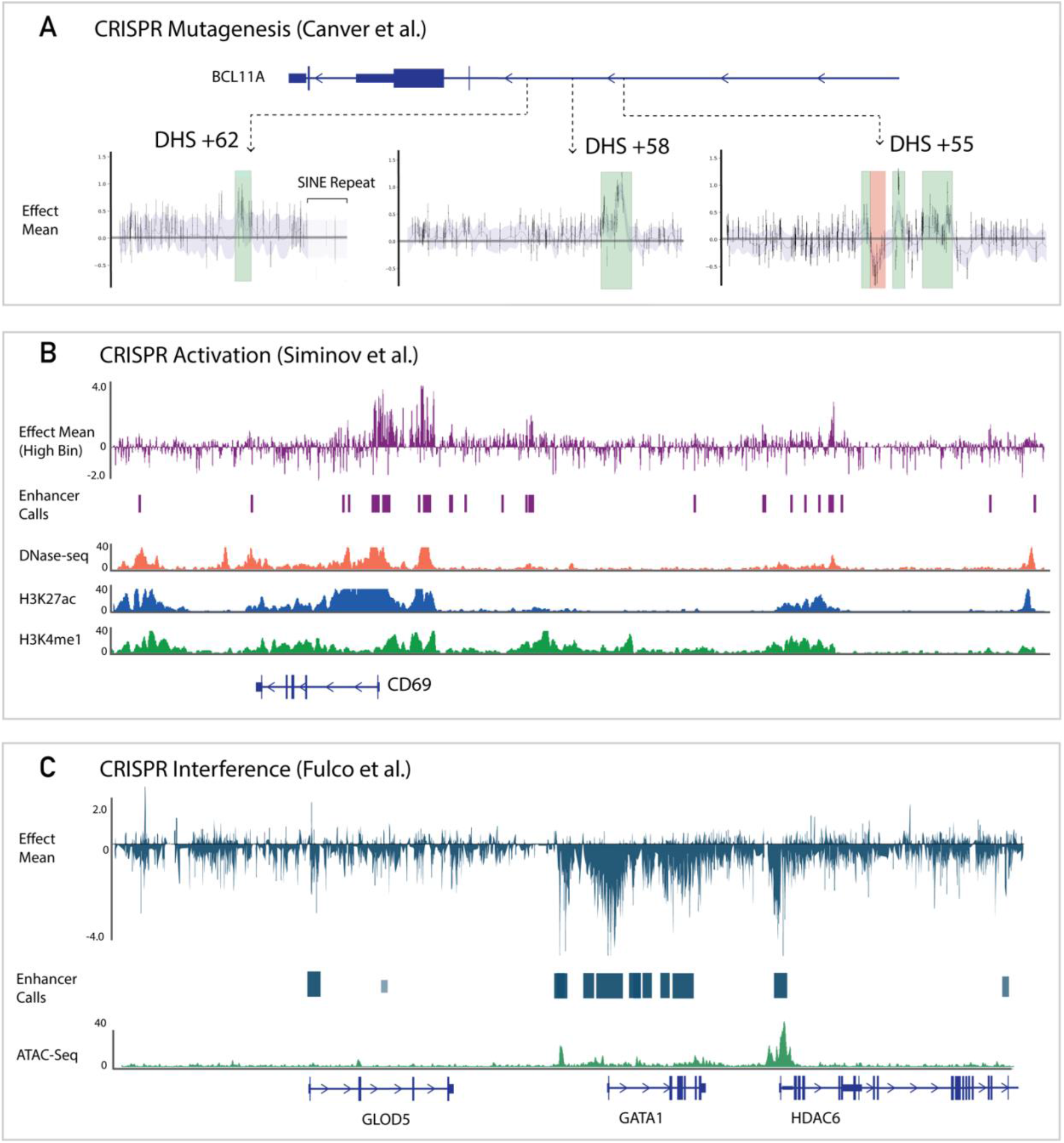
Regulatory elements classified by CRISPR-Decryptr for three published noncoding screens. **A**: Analysis mutagenesis screen targeting BCL11A DHS sites reveals similar enhancer and silencer locations as in the original publications. **B**: Analysis CRISPRa screen targeting CD69 promoter region reveals novel enhancer calls. **C**: Analysis CRISPRi screen targeting GATA1 gene loci reveals the same enhancer calls as in the original analysis.

We have described a statistical technique for analyzing CRISPR noncoding screen data and illustrated the accuracy of CRISPR-Decryptr on three distinct perturbation technologies, demonstrating the method’s ability to reveal novel insights from a diverse set of experimental designs. CRISPR-Decryptr will be a valuable component in future attempts to identify functional genomic elements and their link to phenotypic traits, enabling target identification and synthetic biology in biomedical and biotechnological settings.

## Supplemental Notes

## Section 1: Using CRISPR-Decryptr

In this section, we present the usage and descriptions of each command in CRISPR-Decryptr for noncoding screens. https://github.com/anders-w-rasmussen/crispr_decryptr

### *Step 1:* Infer perturbation-specific regulatory effect from gRNA count data

| Positional Arguments: | Description: |
| --- | --- |
| count_file | Tab-delimited file with columns containing gRNA counts from different samples. The first row should contain sample names corresponding to each column. CRISPR-Decryptr will extract columns from this file according to the sample names included in the design_matrix file. You may include other columns for your own reference (such as gene IDs, etc) but they will be ignored. |
| design_matrix | Tab-delimited file of design matrix for specific experimental design. The first column of the file should contain the samples you are considering, all of which must correspond to an entry in the first row of the count_file. The first row contains the names of the 'effects' you are considering. The rest of the file is 0s and 1s indicating which effects are to be considered in the linear predictor (see section 2.2) |
| replicate_information | This file should have an identical first column to the design matrix. The second contains the word 'replicate' in the first row followed by integers indicating which replicate the sample in the first column is from. |
| Optional Arguments: | Description: |
| --n_chains | Number of chains to use in MCMC sampling (default is 4). |
| --batch_size | Number of guides per batch. Please see notes on the collapsed multinomial in section A.1. Larger batch sizes scale poorly. We recommend the default of 1000. |
| --n_batches | The number of batches to run simultaneously. The number of processes spawned will be n_chains * n_batches. |
| --n_samples | Number of samples for each chain. Default is 500. |
| --outfile | Out filename |
| --sample_file_prefix | Prefix for sample filenames to be saved. The default is "samples." |
| --logfilename | Specify the name for the .log file |
| --outfile_devs | Write replicate level standard deviations of effects to this file. If not specified this file will not be written. These are not used by CRISPR-Decryptr. |

### *Step 2:* Create the convolution matrix

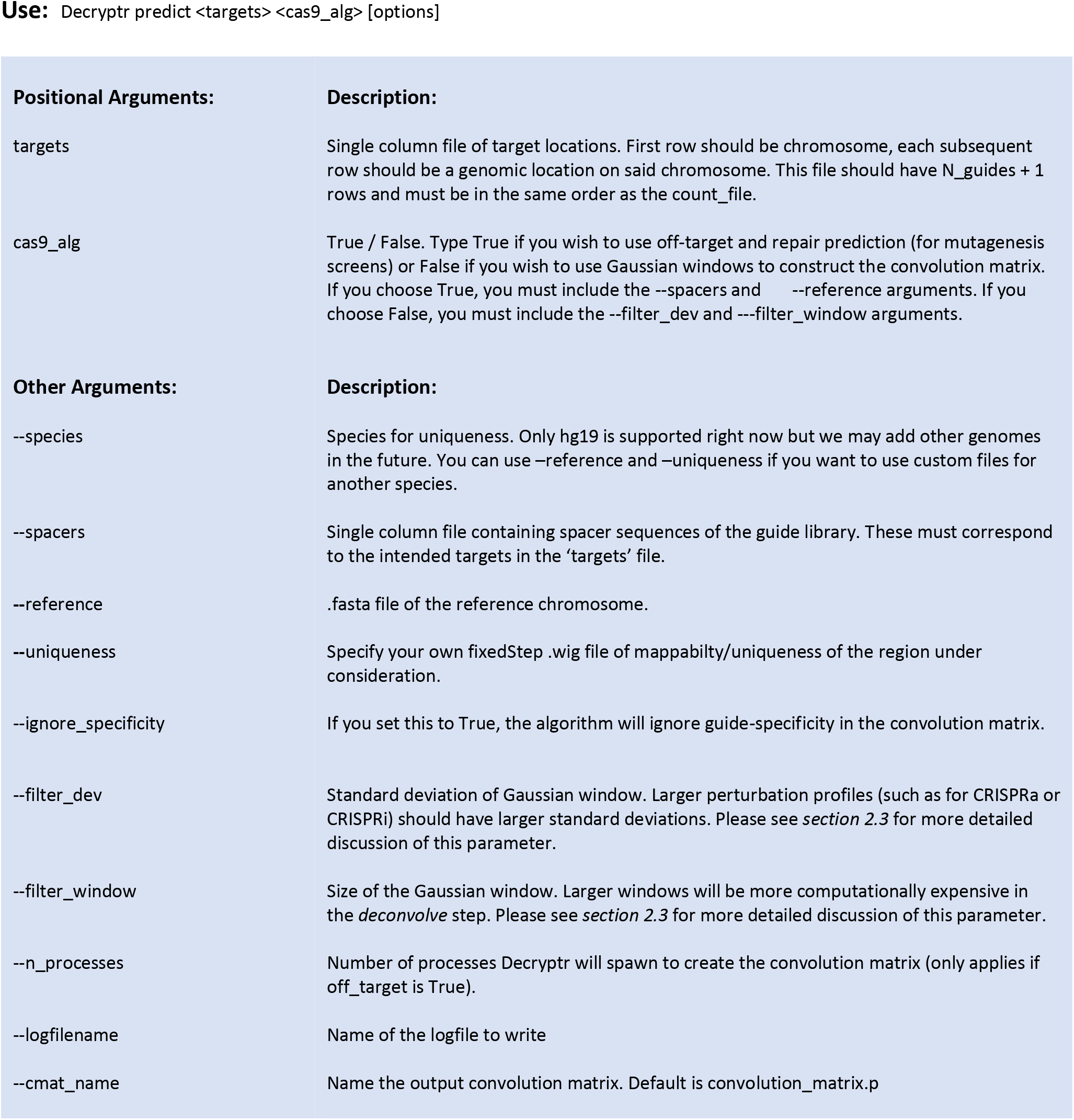

### *Step 3:* Classify regulatory elements with GP deconvolution and HsMMs

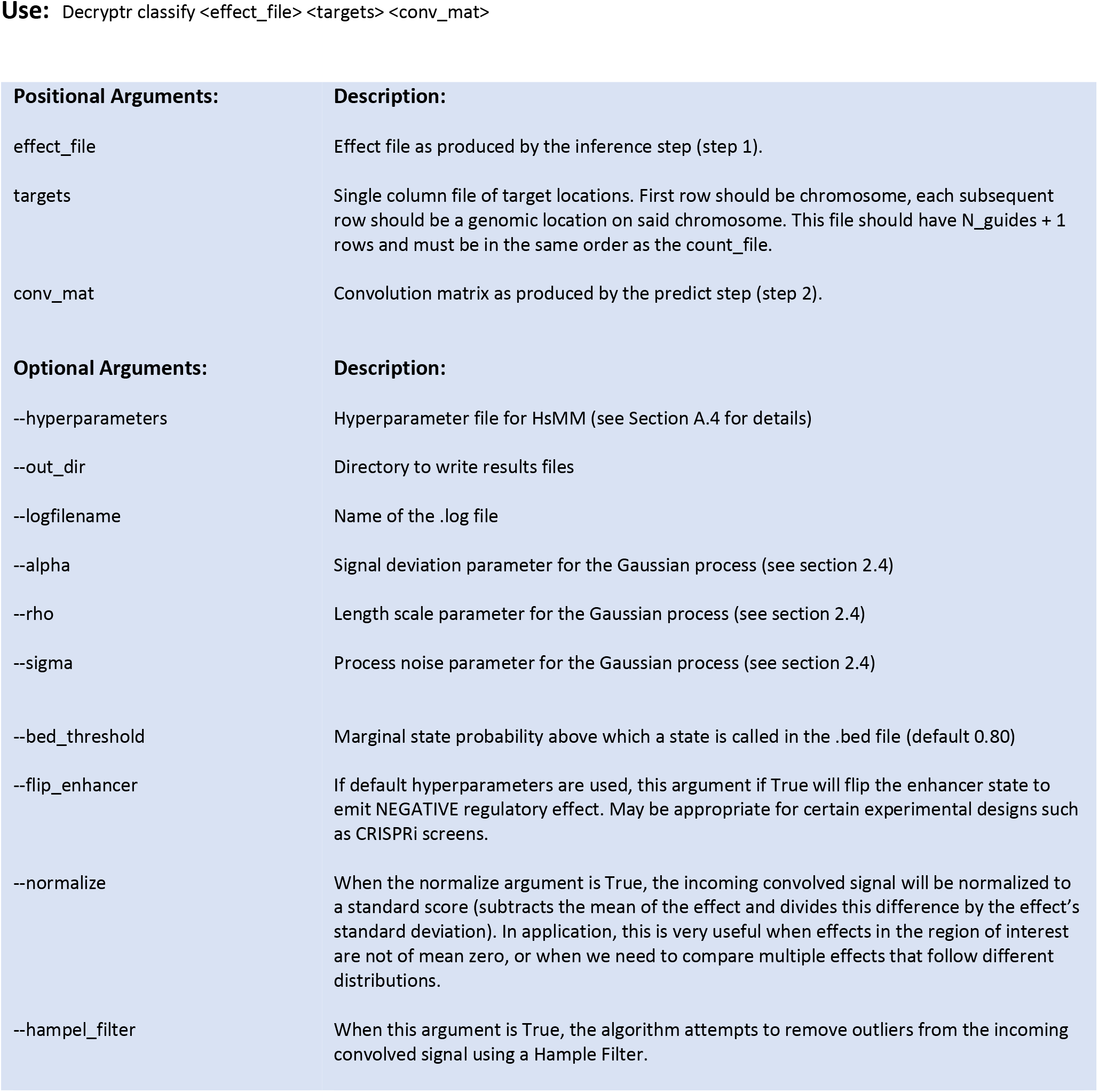

## Section 2: Methods

The objective of CRISPR-Decryptr is to reveal probabilistic locations of regulatory elements on the genome given gRNA counts from noncoding screens. We define a “regulatory landscape,” the set of hidden states at each base of the genomic region of interest (*Section 2.1*). To arrive at these state probabilities, we begin by accounting for experimental design and readout, inferring regulatory effects along the genome using a hierarchical generalized linear model (GLM) (*Section 2.2*). Next, we account for off-target and repair prediction, creating a convolution matrix mapping “guide-specific” effects to “base-specific” effects (Section 2.3). Finally, we implement a Gaussian process (GP) model and hidden semi-Markov model (HsMM) to deconvolve effects and classify the “regulatory landscape” at the base-by-base level (*Section 2.4, 2.5, 2.6*).

### 2.1 Defining the Regulatory Landscape

We assume that perturbation of the noncoding region of interest can have any of *E* user defined effects on phenotype. As such, the probability mass function of phenotypic readout for any perturbation *x*_*pertubation*_ can be written as a function of these effects, *f*_*pertubation*_ (*e*_1_, *e*_2_, ‥ , *e*_*E*_). The goal of the CRISPR-Decryptr method is to take a dataset of phenotypic readouts for various targeted perturbations ***X*** and infer the probabilistic locations of regulatory elements within the targeted region that regulate these *E* effects. We posit that each base *b* ∈ {1, … , *B*} within the genomic region of length *B* belongs to one of *S*_*e*_ states. Denote the state of each *e*, *b* pair *s*_*e*,*b*_. A unique configuration of the entire region across effects is a set of states ***s*** = ⋃_*e ϵ* {1,…,*E*},*b ϵ* {1,…,*B*}_ {*s*_*e*,*b*_}. Given our screen data ***x***, we wish to determine the marginal probability of each (*e*, *b*) pair belonging to any given state, represented by the set of probability vectors

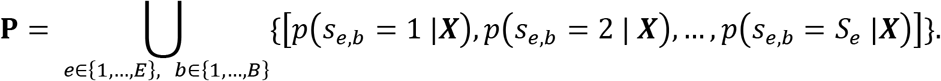

In the following sections, we walk through the model step-by-step, moving from the input data ***X*** to the set of marginal state probability vectors ***P***.

### 2.2 Inferring Regulatory Effect

The first part of our algorithm implements a generalized linear model (GLM) and Markov chain Monte Carlo to infer posterior distributions for effects of each perturbation on cell phenotype, denoted ***g***_*e*_ ∈ ℝ^*I*^ ∀ *e ϵ* {1, … , *E*}, where *I* is the number of guides.^1,2^ Allow *c ϵ* {1, … , *C*} to be condition within the set of *C* conditions, and *r ϵ* {1, … , *R*} to be the replicate within the set of *R* replicates. gRNA count data for each *c*, *r* pair is denoted by the vector 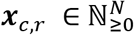 consisting of individual guide specific counts *x*_*c*,*r*,*i*_ where *i ϵ* {1, … , *I*}. We model ***x***_*c*,*r*_ as being generated from a multinomial distribution with parameter ***p***_*c*,*r*_. By using the standard unit *Softmax* function to map a real valued vector in ℝ^*N*^ to the simplex Δ^*N*−1^, we represent ***p***_*c*,*r*_ as a function of a hierarchical linear predictor ***p***_*c*, *r*_ = *Softmax*(***ϕ***_*c*, *r*_). The linear predictor takes the following form:

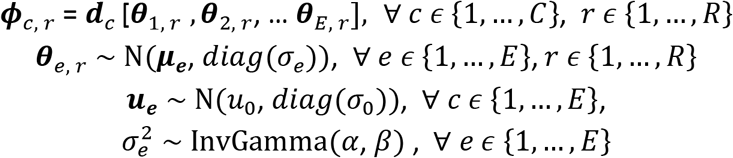

The predictor ***ϕ***_*c*, *r*_ is a linear combination of replicate-specific effect vectors ***θ***_*e*, *r*_ *ϵ* {***θ***_1, *r*_, … , ***θ***_*E*, *r*_} with weights dictated by ***d***_*c*_ row *c* of the design matrix ***D***. Each entry in the replicate-specific effect vector is distributed normally according to *u*_*e*,*i*_ and *σ*_*e*_: guide-specific regulatory effect and a single inferred standard deviation shared across guides. *u*_0_, *σ*_0_, *α*, *β* are fixed hyperparameters. Allow ***ϑ*** = { ***θ***_**1**, **1**_ … , ***θ***_***E***, ***R***,_ ***μ***_**1**_, … , ***μ***_***E***,_ *σ*_1_, … , *σ*_*e*_} to be the set of model parameters and ***ζ*** = {*u*_0_, *σ*_0_, *α*, *β*} to be the set of hyperparameters. We write the joint posterior distribution for the model variables using Bayes formula as,

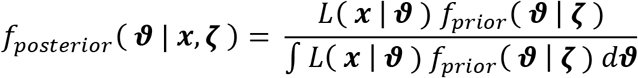

Where *f*_*posterior*_ is the posterior density function of the model parameters given the data and hyperparameters, *f*_*prior*_ is the prior density function of the model parameters given the hyperparameters, and *L* is the likelihood of the data given model parameters. Closed-form representation of the joint posterior is not possible, as evaluation of the marginal likelihood is intractable. To arrive parameter estimates, we sample from the posterior distribution using Hamiltonian Monte Carlo (HMC) implemented in Stan, which will produce *H* sampled values of each parameter. Allow ***ϑ***_*h*_ to be the set of sampled parameters for sample *h ϵ* {1, … , *H*}. We calculate the posterior means of each ***μ***_*e*_, and 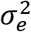 as follows:

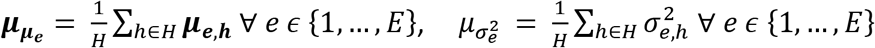

We can now express the guide-specific regulatory effects as,

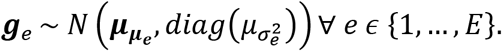

**Supplementary Figure 2.2:**
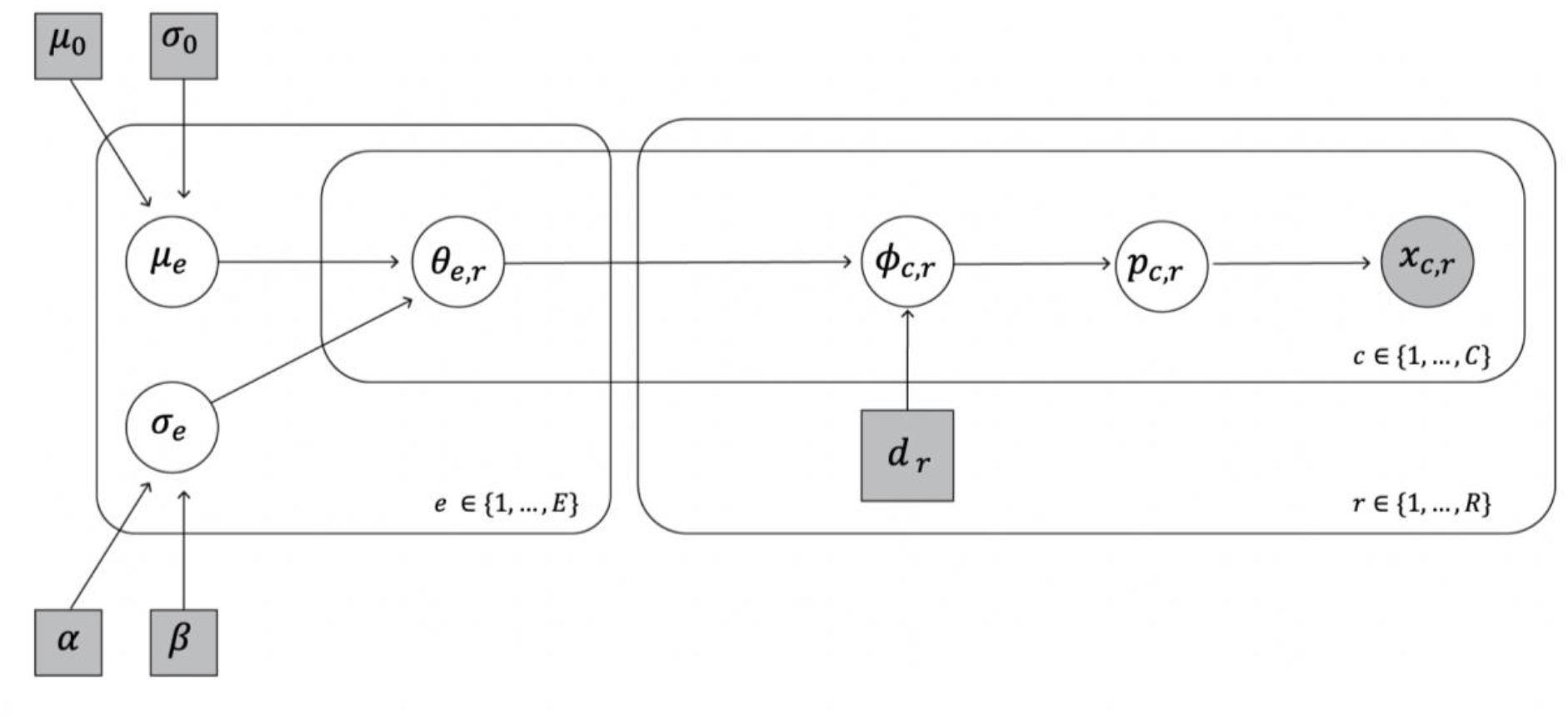
Plate diagram of hierarchical Generalized Linear Model for effect inference. White circles represent latent model parameters to be inferred, while grey circles represent observed quantities. Grey squares are user-defined parameters, while circular nodes are model variables.

### 2.3 Constructing a Convolution Matrix

In *Section 2.1*, we defined the latent regulatory landscape ***s*** as being the set of base and effect-specific states ⋃_*e ϵ* {1,…,*E*},*b ϵ* {1,…,*B*}_ {*s*_*e*,*b*_}. In *Section 2.2* above, we described a numerical method to infer perturbation specific effects ***g***_*e*_ ∈ ℝ^*I*^. However, to examine regulatory elements at the base-by-base level of granularity we must arrive at a base-specific effect vectors ***b***_*e*_ ∈ ℝ^*B*^ ∀ *e* ∈ {1, … , *E*}. We do so by positing that there exists a linear map *f*: ℝ^*B*^ → ℝ^*B*^ that can be represented as an *I* × *B* convolution matrix ***C***, such that ***g***_*e*_ = ***Cb***_*e*_ ∀ *e* ∈ {1, … , *E*}. We define the entries *c*_*ib*_ of matrix ***C*** as representing the probability of guide *i* perturbing base *b*. As such, we construct the convolution matrix guide-by-guide by calculating Cas9-binding and perturbation probabilities across the region of interest. The raw data, as well as our inferred effects, are specific to guide *i* with target sequence, *target*_*i*_. We define the target DNA sequence as the sequence complementary to the guide RNA. For mutagenesis screens, we assume the probability of guide *i* inducing a mutation at base *b* can be written

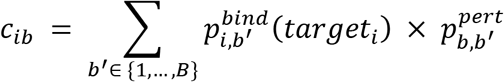

where *b*′ is the base where Cas9 binds, defined as the most 3’ nucleotide in the PAM sequence NGG on the both strands. 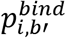 is the probability that Cas9 fused with guide *i* binds at base *b*′. 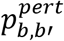 is the probability that a Cas9 binding event at base *b*′ results in a genomic perturbation at base *b*.

We calculate each binding probability 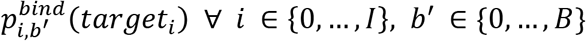 using the scoring methodology presented in Hsu *et al.* (2013)^3^. Any base that is not the last nucleotide in an NGG motif is assigned a Cas9 binding of zero. Allow *target*_*i*_[*b*] to be the b-th nucleotide in the target sequence and *candidate*_*b*′_[*b*] to be the b-th nucleotide in the 20 base-pair sequence beginning 23 bases in the 5’ direction of *b*′ and terminating 3 bases in the 5’ direction of *b*′. We define the Hsu tensor *Hsu*[*t*, *c*, *b*] as a 4 × 4 × 20 tensor of Hsu scores defined in **Supplementary Table 2.3**.

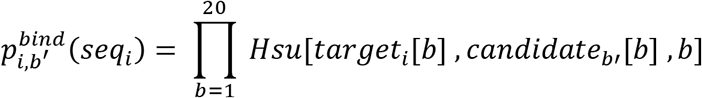

The prediction of perturbation profile is dependent on experimental design, for CRISPR mutagenesis screens that induce cleavage events, we can calculate perturbation probabilities at each base around the binding site 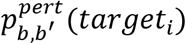 using a Convolutional Neural Network trained on a database of double stranded break outcomes in Allen et al (2017)^4^. Please see *Section A.3* on more details on CNN model performance. Allow ***seq***_***b***′_ to be a 4 × 41 one-hot encoding of the sequence beginning 20bps in the 5’ direction of *b*′ and ending 20bps in the 3’ direction of *b*′. We allow 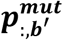 to be the 1 × 41 vector 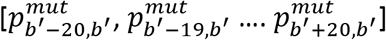. As such, the CNN model approximates a function 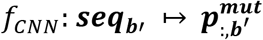.

**Supplementary Figure 2.3:**
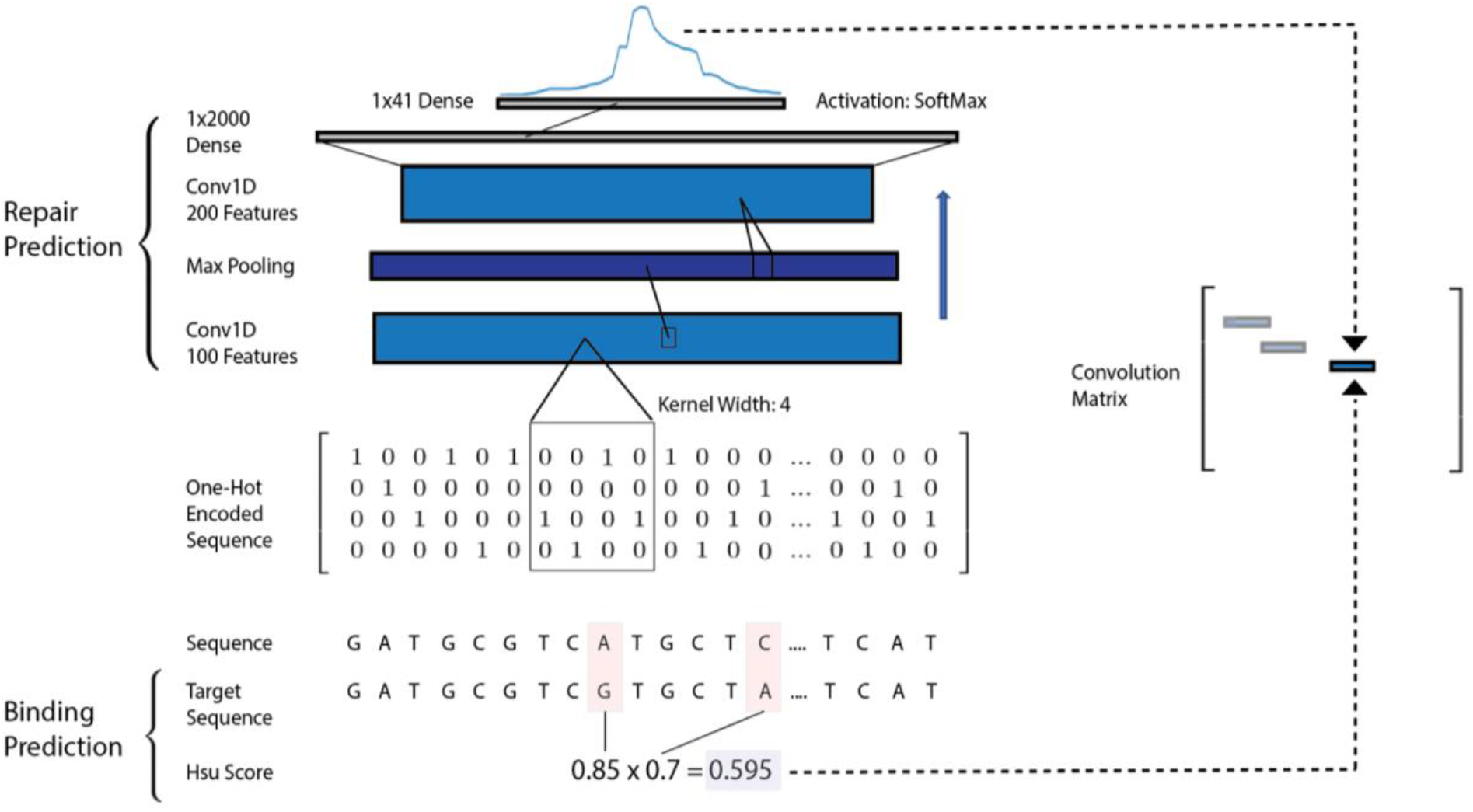
Illustration CRISPR-Decryptr’s method for convolution matrix construction. For Cas9 mutagenesis screens, repair prediction is performed by a convolutional neural network model, taking one-hot encoded sequence as an input and outputting a sequence-specific pertubation profile. The guide is also aligned to the reference genome based on its Hsu score, determining the position and weighting the perturbation profile in the convolution matrix.

For CRISPRi and CRISPRa screens, we construct the convolution matrix out of Gaussian Windows of user defined standard deviation *σ*_*window*_ and odd integer window size *W*, centered on *b*′_*i*_, defined as the most 3’ nucleotide in the PAM sequence of the intended target of guide *i*. We write the entries of the convolution matrix as follows:

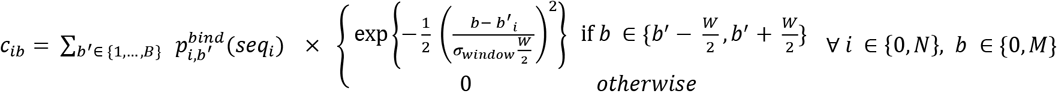

**Supplementary Table 2:** Illustration of the CRISPR-Decryptr method for convolution matrix construction. For Cas9 mutagenesis screens, repair prediction is performed by a convolutional neural network model, taking one-hot encoded sequence as an input and outputting a sequence-specific pertubation profile. The guide is also aligned to the reference genome based on its Hsu score, determining the position and weighting the perturbation profile in the convolution matrix.

|  | 19 | 18 | 17 | 16 | 15 | 14 | 13 | 12 | 11 | 10 | 9 | 8 | 7 | 6 | 5 | 4 | 3 | 2 | 1 |
| --- | --- | --- | --- | --- | --- | --- | --- | --- | --- | --- | --- | --- | --- | --- | --- | --- | --- | --- | --- |
| A:A | 1.19 | 1.10 | 2.87 | 1.13 | 0.52 | 0.64 | 1.03 | 0.54 | 1.29 | 0.08 | 0.26 | 0.27 | 0.05 | 0.03 | 0.01 | 0.02 | 0.17 | 0.70 | 0.27 |
| A:C | 1.51 | 1.16 | 1.66 | 0.99 | 0.02 | 0.55 | 1.87 | 0.77 | 0.93 | 0.73 | 0.48 | 0.48 | 0.71 | 1.02 | 0.02 | 0.35 | 0.18 | 0.14 | 0.27 |
| A:G | 1.48 | 1.08 | 1.30 | 0.74 | 0.98 | 0.49 | 0.93 | 0.60 | 1.04 | 0.30 | 0.82 | 0.03 | 0.04 | 0.06 | 0.03 | 0.16 | 0.10 | 0.22 | 0.03 |
| A:T | 1.00 | 1.00 | 1.00 | 1.00 | 1.00 | 1.00 | 1.00 | 1.00 | 1.00 | 1.00 | 1.00 | 1.00 | 1.00 | 1.00 | 1.00 | 1.00 | 1.00 | 1.00 | 1.00 |
| C:A | 1.02 | 0.96 | 2.25 | 1.62 | 0.24 | 0.76 | 1.07 | 0.11 | 1.43 | 0.35 | 0.62 | 0.73 | 0.12 | 0.04 | 0.50 | 0.41 | 0.18 | 0.53 | 0.17 |
| C:C | 0.03 | 1.05 | 1.05 | 1.41 | 0.05 | 0.64 | 2.08 | 0.46 | 0.16 | 0.14 | 0.10 | 0.04 | 0.01 | 0.05 | 0.01 | 0.21 | 0.06 | 0.03 | 0.06 |
| C:G | 1.00 | 1.00 | 1.00 | 1.00 | 1.00 | 1.00 | 1.00 | 1.00 | 1.00 | 1.00 | 1.00 | 1.00 | 1.00 | 1.00 | 1.00 | 1.00 | 1.00 | 1.00 | 1.00 |
| C:T | 1.67 | 0.97 | 1.07 | 1.09 | 0.65 | 1.01 | 0.86 | 0.31 | 0.45 | 0.90 | 0.43 | 0.56 | 0.14 | 0.07 | 0.33 | 0.24 | 0.04 | 0.14 | 0.44 |
| G:A | 1.17 | 0.45 | 2.79 | 1.05 | 0.60 | 0.81 | 0.79 | 0.69 | 1.37 | 0.12 | 0.20 | 0.25 | 0.01 | 0.02 | 0.05 | 0.37 | 0.32 | 0.56 | 0.46 |
| G:C | 1.00 | 1.00 | 1.00 | 1.00 | 1.00 | 1.00 | 1.00 | 1.00 | 1.00 | 1.00 | 1.00 | 1.00 | 1.00 | 1.00 | 1.00 | 1.00 | 1.00 | 1.00 | 1.00 |
| G:G | 1.32 | 1.42 | 1.30 | 1.21 | 0.95 | 0.23 | 1.02 | 0.81 | 1.16 | 0.71 | 1.58 | 0.35 | 0.09 | 0.05 | 0.00 | 0.23 | 0.22 | 0.53 | 0.50 |
| G:T | 1.65 | 0.90 | 1.60 | 0.92 | 0.81 | 0.96 | 1.02 | 0.27 | 0.71 | 1.52 | 0.69 | 1.04 | 0.53 | 0.90 | 0.72 | 0.84 | 0.54 | 0.28 | 0.69 |
| U:A | 1.00 | 1.00 | 1.00 | 1.00 | 1.00 | 1.00 | 1.00 | 1.00 | 1.00 | 1.00 | 1.00 | 1.00 | 1.00 | 1.00 | 1.00 | 1.00 | 1.00 | 1.00 | 1.00 |
| U:C | 0.04 | 1.00 | 1.65 | 1.34 | 0.01 | 0.54 | 2.34 | 0.44 | 0.21 | 0.34 | 0.26 | 0.09 | 0.05 | 0.07 | 0.02 | 0.37 | 0.05 | 0.09 | 0.25 |
| U:G | 1.54 | 1.28 | 1.27 | 1.22 | 1.06 | 0.78 | 1.24 | 0.99 | 1.53 | 0.72 | 1.74 | 0.42 | 0.13 | 0.07 | 0.05 | 0.23 | 0.31 | 0.13 | 0.06 |
| U:T | 1.73 | 0.82 | 1.16 | 1.18 | 0.74 | 0.90 | 0.84 | 0.30 | 0.62 | 1.14 | 0.36 | 0.62 | 0.18 | 0.27 | 0.28 | 0.26 | 0.12 | 0.17 | 0.34 |

### 2.4 Gaussian Process Deconvolution

In *section 2.3* we constructed a convolution matrix that satisfies the equation ***g***_*e*_ = ***Cb***_*e*_ under our assumptions. However, the problem of solving for a vector of base-specific effect ***b***_*e*_, given a vector of guide-specific effect, is ill-posed, as the matrix ***C*** is singular. To solve the inverse problem, we implement a Gaussian Process model by defining a Gaussian process prior on ***b***_*e*_ such that ***b***_*e*_ ~ *GP*(**0**, ***K***)^5^. ***K*** is a kernel matrix approximating the covariance structure as a squared exponential function of the distance between guides *i* and *j*, 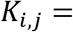 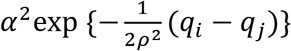, where *q*_*i*_, *q*_*j*_ are the genomic positions of *i* and *j* respectively. The characteristic-length scale *ρ* and signal variance *α*^2^ are inferred parameters. Note that ∘ denotes the Hadamard or element-wise product.

Given the linear map ***C*** defined in 2.3, we write the likelihood for the guide specific effect variables as ***g***_*e*_ ~ ***N***(***Cb***_*e*_, *diag*(*σ*^2^)) where *σ*^2^ is a process-noise term to be inferred. The joint distribution of ***g***_*e*_ and ***b***_*e*_ can now be written:

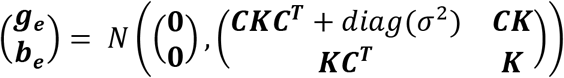

The posterior predictive distribution for ***b***_*e*_ now takes the closed-form solution,

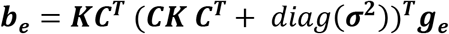

The parameters *ρ*, *α*^2^, *σ*^2^ can optionally be set by the user. By default, CRISPR-Decryptr sets *α*^2^ and *σ*^2^ both to the average variance in the guide-specific effect signal and sets the length scale *ρ* to the average distance between saturated guides (defined as guides within 100bps).

By default, CRISPR-Decryptr does not normalize the incoming effect ***g***_*e*_. However, if desired, the classify step can use the convolved effect’s standard score 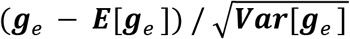 if the --normalize argument is set to True (see *section 1*). As the GP prior has a fixed mean of zero, this can influence the results of the deconvolution if the difference between zero and ***E***[***g***_*e*_] is large.

### 2.5 Decoding Latent State Variables with Hidden semi-Markov Models

We model the latent state path as following a discrete time semi-Markov process, where the effect specific state of each base is denoted *s*_*e*,*b*_. In a standard Markov process, the duration – or sojourn time – spent within any given state, follows a geometric distribution, an assumption we believe to be in conflict with the distribution in sizes of experimentally measured regulatory elements. As such, we implement a Hidden semi-Markov Model with state durations that follow a negative binomial distribution with state-specific parameters, written *D*_*s*,*e*_ ~ *NB*(*r*_*s*,*e*_, *p*_*s*,*e*_)^6^. For CRISPR-Decryptr, we assume the HsMM has a latent model structure comprised of an arbitrary number of states *n* (default is two). The transition matrix and initial probability vector for each effect *e* take the form:

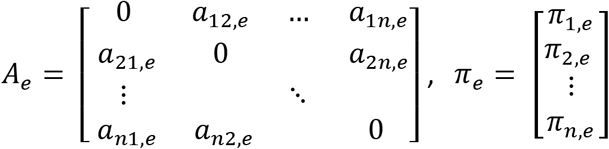

Where *a*_*ij*,*e*_ represents the probability of transitioning from effect-specific state *i* to state *j* and *π*_*i*,*e*_ represents the probability of base 0 being the effect-specific state *i*.

We now define the emission probabilities for each state. We assume the base-specific mean for effect *e*, *μ*_*e*,*b*_ follows a Normal distribution with inferred mean *μ*_*e*,*s*_ and precision 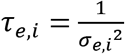, where *σ*_*e*,*b*_ was the standard deviation of the effect mean at base *b.* We write this probability as follows:

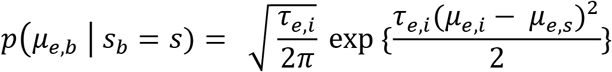

To maintain a Bayesian approach, we define prior distributions on model parameters as follows:

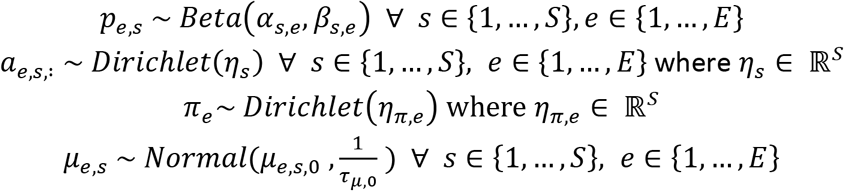

Implementation of the Baum-Welch algorithm leads to estimates for marginal state-probability vectors ***P*** as described in *section 2.1.*

## Section 3: Analysis of Published Screens

### 3.1 Mutagenesis Screen of BCL11A DNase Hypersensitivity Sites (Canver et al.)

The BCL11A gene is a validated repressor of fetal hemoglobin (HbF) level. In their 2013 paper, Bauer et al. label three DNAse hypersensitivity sites (DHS) that overlap with single nucleotide polymorphisms (SNPs) that impact HbF levels in genome-wide association studies (GWAS)^7^. These sites, DHS +62, +58, and +55, are named according to their distance in kilobases from the TSS of BCL11a. The HbF-associated SNPs rs1427407 and rs7606173 are located within DHS +62 and DHS +55 respectively. DHS +58 has proven a successful therapeutic target in patients with hemoglobinopathies using CRISPR-Cas9 gene editing technology. Due to their function being known *a priori* and validated in application through therapeutic interventions, these three sites are ideal for exploring experimental and computational approaches to interrogating cis-regulatory elements. In Canver et al., these sites are perturbed in human umbilical cord blood-derived erythroid progenitor (HUDEP) cells CRISPR-Cas9^8^. The phenotypic readout for this experiment is gRNA counts from cells FACS sorted into a HbF high and HbF low bin and an early condition as control. The original analysis defines HbF enrichment as the log_2_ fold-change between the normalized HbF high and HbF low bins and utilizes a three-state Hidden Markov Model (HMM) to classify silencers and enhancers. We re-analyze this data with CRISPR-Decryptr, implementing a design matrix for effect inference as follows:

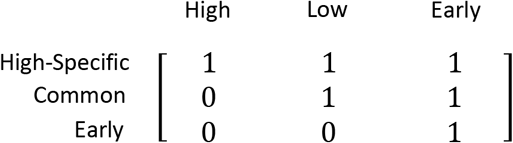

We utilize the following hyperparameters for the Hidden semi-Markov Model:

| STATE | E_MU | E_TAU_0 | NB_R | NB_SUCC | NB_FAIL |
| --- | --- | --- | --- | --- | --- |
| BACKGROUND | 0 | 1 | 2 | 1 | 500 |
| SILENCER | -2 | 1 | 2 | 1 | 40 |
| ENHANCER | 2 | 1 | 2 | 1 | 40 |

From the high-specific effect, we arrive at a regulatory landscape in concordance with the original analysis. The common and early effects also yield insight into other phenomena impacting gRNA counts in the other bins. Targeting the Alu SINE repeat in the DHS +62 region impacts cell viability in both the early and later bins, a phenomenon that was also noted in the original analysis.

**Supplementary Figure 3.1.1:**
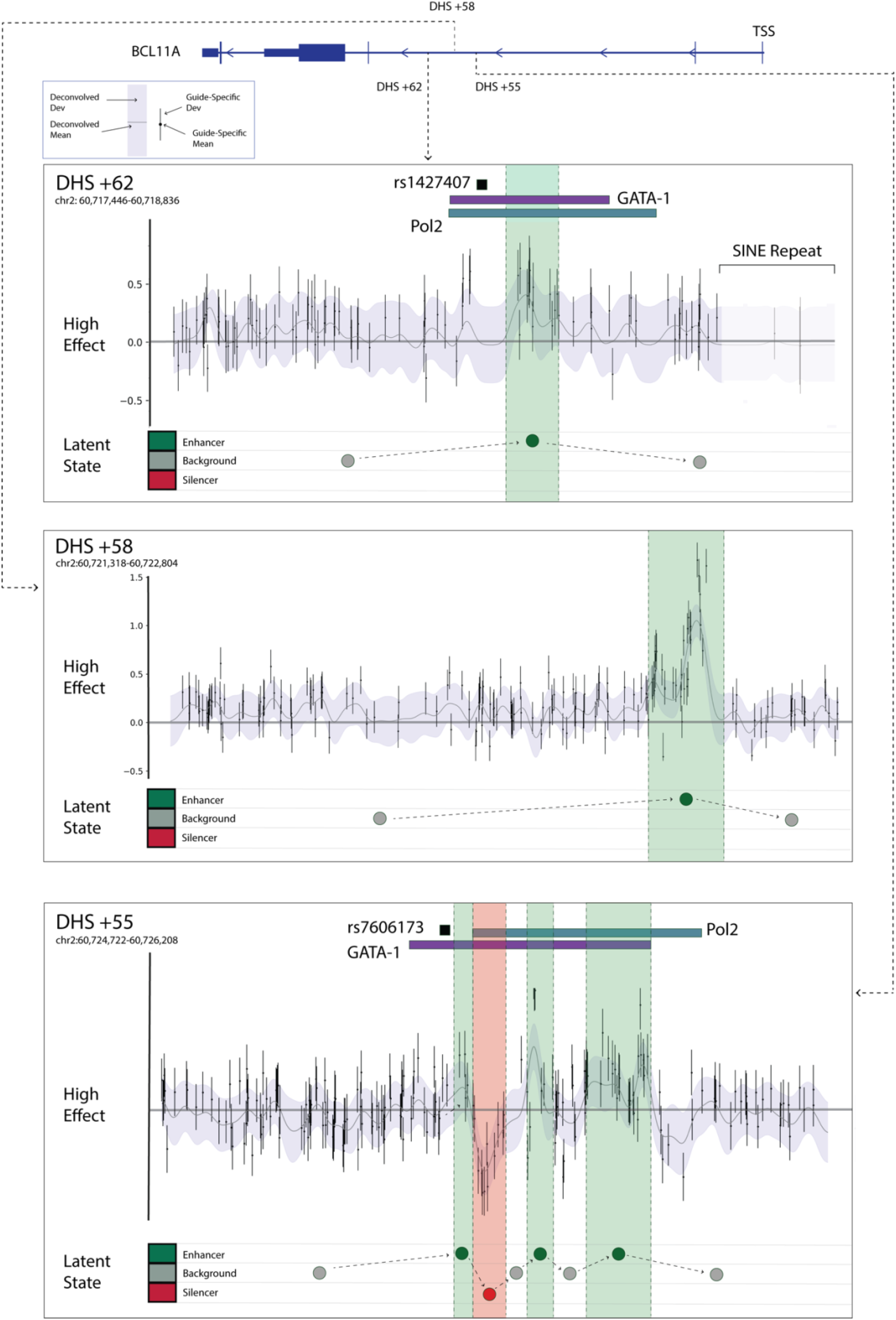
Results from the analysis of guides targeting BCL11A DHS hypersensitivity sites in Canver et al. The high-specific effect reveals similar enhancer and silencer calls as in the original analysis.

### 3.2 CRISPR Activation Screen at IL2RA and CD69 Gene Loci (Simeonov et al.)

Simeonov et al. introduce the CRISPR activation screen, which utilizes a mutated version of Cas9 (dCas9) without endonuclease activity which - when fused with gRNA and transcriptional activators – is able to activate regulatory elements on the genome^9^. To validate the approach, the authors target IL2RA and CD69 gene loci in two different pooled screens, by transducing Jurkat-T cells with a dCas9-VP64 activator. To measure phenotypic change, the authors FACS sort cells into a “negative”, “low”, “medium”, and “high” bins of IL2RA and CD69 based on expression levels, as well as a non-transduced control group. We re-analyze this screen using the following design matrix:

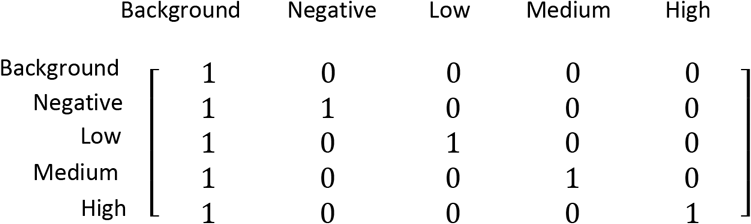

For the creation of the convolution matrix, we use the gaussian window method described in *section 2.3* with default parameters. For the HsMM, we utilized default hyperparameters.

CRISPR-Decryptr successfully recalls all the putative regulatory elements from the Simionov et al. analysis. The two putative regulators CaRE1 and CaRE6, revealed solely in the “low” bin in the original analysis, are now revealed “medium” bin as well. Additionally, CRISPR-Decryptr calls novel putative enhancers of varying strengths, many of which are highly correlated with DNAse-seq and the active enhancer mark H3K27ac from the Jurkat-T Cell line. These putative enhancers include the nearby RBM17 and PFKFB3 gene promoters.

Putative enhancers of different strengths (marginal state probability > 0.95, > 0.85, and > 0.75 for Strong, Moderate, and Faint calls respectively) included in **figure 3.2.2** illustrates how the method’s output of marginal state probabilities provides an easily interpretable metric by which to rank regulators.

**Figure 3.2.2:**
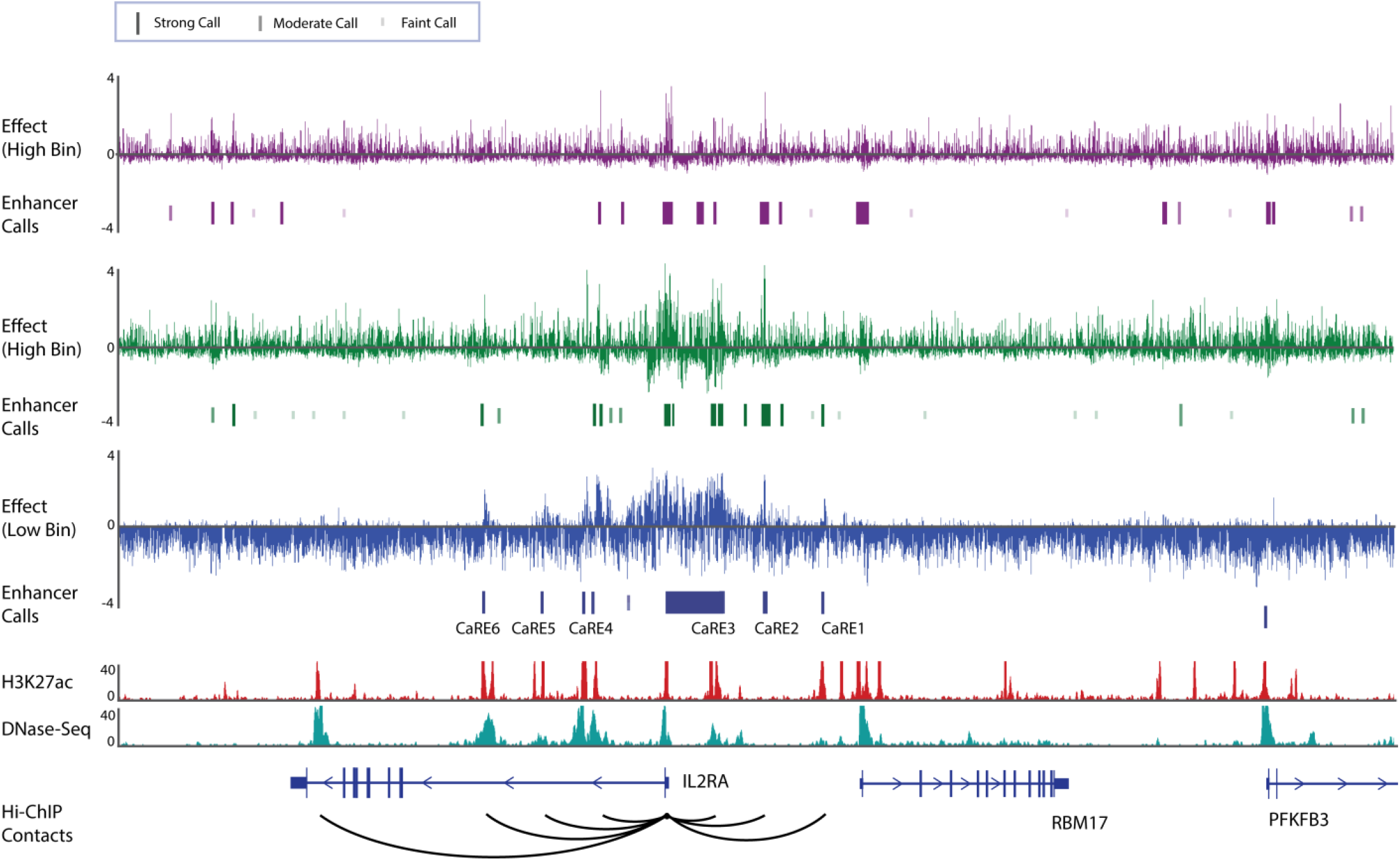
Analysis of Simenov et al. CRISPRa screen of the IL2RA gene locus. High, Medium, and Low effects are inferred from the inference step of CRISPR-Decryptr, with enhancer state calls from the HsMM of varying strengths displayed below respective bins (marginal state probability > 0.95, > 0.85, and > 0.75 for Strong, Moderate, and Faint calls respectively). H3K27ac, DNAse-seq, and HiChIP contacts, are correlated with active enhancers.

At the CD69 gene loci, we consider the “high” expression bin as in the original analysis. Similar to the IL2RA gene loci, we recall the original enhancers and elucidate new enhancer calls in strong agreement with chromatin accessibility data and histone marks.

**Figure 3.2.3:**
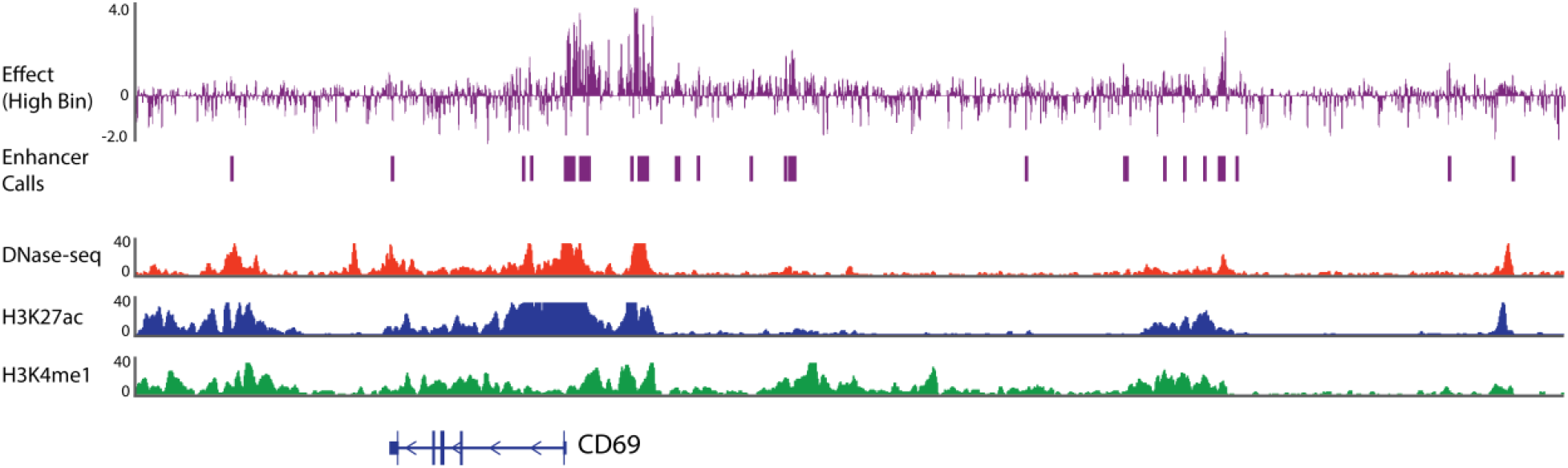
Analysis of Simenov et al. CRISPRa screen of the CD69 gene locus with the deconvolved effect from the “high” bin displayed above “strong” enhancer calls (p > 0.95). H3K27ac, DNAse-seq, and H3K4me1 are displayed to compare results with chromatin accessibility and active enhancer markers.

### 3.3 CRISPR Interference Screen at the GATA1 Gene Locus (Fulco et al.)

In a noncoding screen targeting the GATA1 loci, Fulco et al. fuse inactivated dCas9 with Krüppel associated box (KRAB) domain to repress transcriptional activation – an approach termed CRISPR Interference^10^. K562 erythroleukemia cells which express the KRAB-dCas9 doxycycline-inducible promoter were infected with a custom gRNA library. As GATA1 impacts K562 proliferation, phenotypic readout is gRNA counts, where depletion implies decreased gene expression. As such, the targeting of an enhancer should result in negative inferred regulatory effect around the site of perturbation. For the analysis, we utilize a filter standard deviation of 30, set the *--flip_enhancer* argument to True in the classify step, and use a design matrix of the following form:

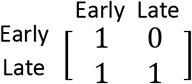

In addition to calling the same putative regulators as in the original analysis, CRISPR-Decryptr reveals one faint novel enhancer at the GATA1 loci within the GLOD5 intron. From ChIPseq of K562 cells, we see this call is at the binding site of a CEBPB transcription factor, a known regulator of inflammatory processes which is also bound at other regions demonstrating gRNA depletion in the GATA1 loci.

**Figure 4.3.1:**
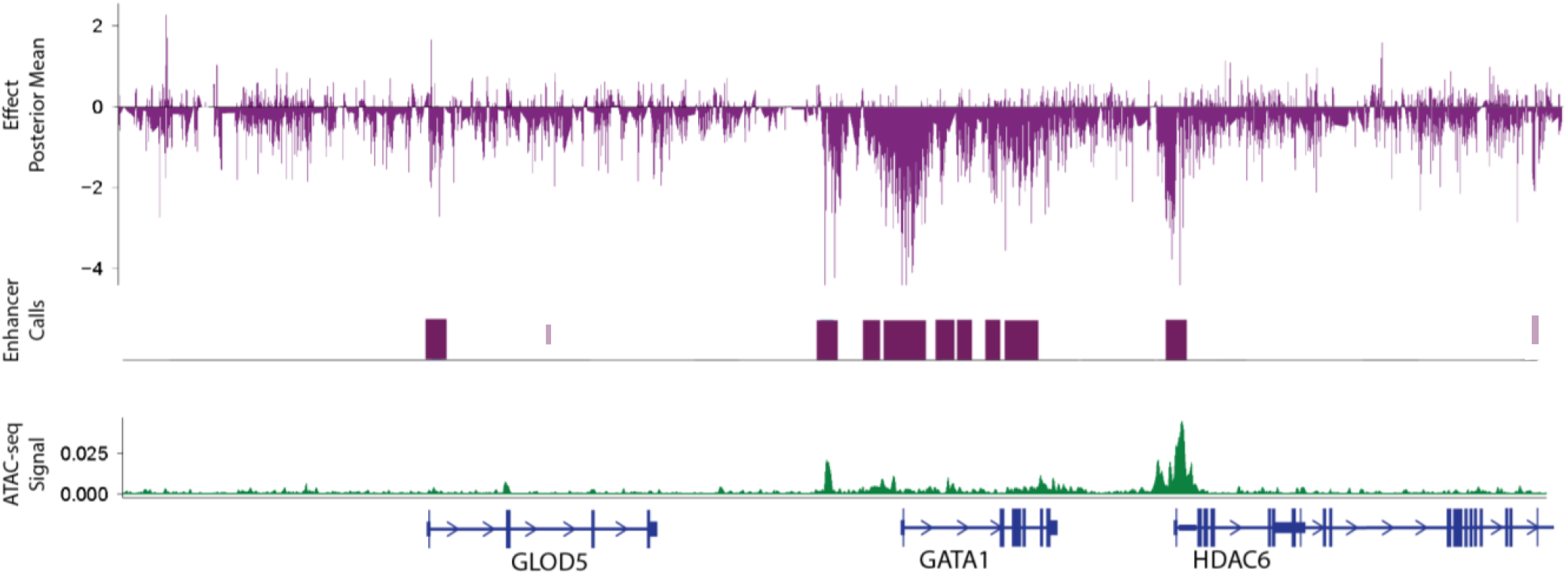
Analysis of gRNAs fused with KRAB-dCas9 targeting the GATA1 gene loci. In addition to enhancers identifies in the original analysis, CRISPR-Decryptr reveals a novel enhancer targeting an experimentally validated CEBPB transcription factor binding site in K562 cells.

## Section 4: Discussion

### 4.1 Discussion of Existing Methods

CRISPR noncoding screens represent a relatively new method for interrogation of functional elements in the noncoding genome. We believe any method aiming to classify *de novo* regulatory elements must be able to accomplish two tasks: firstly, the algorithm should be able to take raw screen data and translate it into some quantity representing the effect of perturbation on phenotype. Second, the algorithm should classify regulatory element locations using this quantity. At the time of this submission, two methods have been published in the literature, both of which address some part of a complete noncoding screen analysis. CRISPR-SURF,^11^ a deconvolution framework employing Lasso regularization, and MAUDE^13^ a method for the analysis of gene expression changes in sorting-based CRIPSR screens, such as the CRISPRa screens discussed in *section 3.2*.

In CRISPR-SURF each gRNA’s effect on phenotype is calculated as the Log_2_ fold change (Log_2_FC) across a pair of conditions and/or timepoints for a specific perturbation. When calculating the ratio between counts, small denominators lead to exaggerated fold changes. Similarly, large count differences have the tendency to be underexaggerated by the transformation. Please see *section A.1.1* for simulated illustrations of the drawbacks of fold changes when applied to simulated gRNA count data. Additionally, the fold change calculation provides no intuitive way to account for experimental design or combine information across replicates without crude calculations such as averaging fold changes. CRISPR-SURF primarily serves as a deconvolution method, modeling experiment specific perturbation profiles and mapping guide-specific quantities to base-specific quantities using a deconvolution operation implemented through Lasso, a common technique in machine learning for solving ill-posed inverse problems. In the case of CRISPR mutagenesis screens, CRISPR-SURF applies a user-defined perturbation profile to all guides, however, research on measured repair outcomes from Cas9 cleavage in Allen et. al. indicates that perturbation profile is sequence specific. Additionally, CRISPR-SURF does not account for off-target effects or sequence specificity, a major concern with any experiment or therapy using CRISPR technology. CRISPR-Decryptr explicitly models sequence specific repair profiles and off-target effects for CRISPR mutagenesis screens, predicting posterior effect variables at each base in the noncoding region using a Gaussian Process (GP) Model for deconvolution. In contrast with techniques such as Lasso or Tikhonov regularization, the GP framework is able to account for uncertainty estimates from the guide-specific inference step when predicting base-specific effect parameters.

MAUDE takes a statistical approach that is designed specifically for binned count readouts, an experimental design where mutated cells are FACS sorted based on their expression level as in the IL2RA/CD69 CRISPRa screen (Simionov et al) presented in *section 3.2*. The authors propose a model that accounts for gRNA count distributions across these FACS sorted bins, accounting for control guides to arrive at guide-specific Z-score which serves as their metric of effect on expression level. As this method is designed specifically to account for binned gRNA counts, it does not have the ability to account for other screen designs. Additionally, the statistical model does not provide a statistical treatment of parameter uncertainty that utilizes information across replicates. For classification of regulatory elements, MAUDE utilizes a sliding window method of arbitrary size to group guides, CRISPR-Decryptr’s HsMM step decodes the marginal probabilities of regulatory state-space configurations of individual nucleotides, fully capturing spatial information in the deconvolved effect signal. In *section A.1.2*, we present simulations to demonstrate the limitations of sliding window analysis and relative outperformance of HsMMs.

Neither CRISPR-SURF nor MAUDE account for guide specificity, off-target effect, or repair outcome. We believe these are factors that should not be ignored when analyzing CRISPR noncoding screens. The authors of MAUDE claim their algorithm outperforms other methods due to the fact that it classifies 12 new regulatory elements when analyzing the CD69 CRISPRa screen (section 3.2). While 10 of these region appear to downregulate CD69, the authors do not account for the fact that these regions are overlapping extremely non-unique sequences, such as SINE repeats. As we discuss in a brief note in *section A5*, non-specific guides appear to be consistently depleted in some experimental designs, including the CD69 screen. By not accounting for guide-specificity, we believe the majority of MAUDEs regulatory element calls in their validation to be false positives. This highlights the importance of considering guide-specificity through all steps of the analysis of CRISPR noncoding screens.

CRISPR-Decryptr is the most complete framework for analyzing CRISPR noncoding screens of various experimental designs. As a fully generative model, CRISPR-Decryptr has an explicit model formulation and clear mathematical assumptions. The Generalized Linear Model implemented in *section 2.2* captures the compositional nature of gRNA counts, and able to model a diverse set of experimental designs through the user defined design matrix. The Gaussian Process utilized in *section 2.4* provides deconvolution framework similar to that in CRISPR-SURF, while also accounting for off-target effects, guide specificity, and probabilistic repair outcomes. The Hidden semi-Markov Model in CRISPR-Decryptr provides accurate state classification at the base-by-base level of granularity. Finally, by employing Bayesian inference techniques, CRISPR-Decryptr models parameter uncertainty at all stages of the analysis.

**Table 4.1.1:** Table comparing how the three algorithms approach what we believe to be the three requisite aspects of a complete CRISPR noncoding screen analysis.

|  | CRISPR-SURF | MAUDE | CRISPR-DECRYPTR |
| --- | --- | --- | --- |
| MODEL FOR INFERENCE OF GUIDE-SPECIFIC EFFECT | Log2 Fold Change | Statistical Model for Binned gRNA Counts | Generalized Linear Model with Flexible Linear Predictor. MCMC Bayesian Inference. |
| MAPPING GUIDE-SPECIFIC EFFECT TO BASE-SPECIFIC EFFECT | Lasso-based Deconvolution | None | Gaussian Process Model with Off-Target Alignment, Specificity Adjustment, and Repair Outcome Prediction |
| CLASSIFICATION METHOD | None | Sliding Window Statistical Tests | Bayesian Hidden semi-Markov Model |

**Table 4.1.2:**
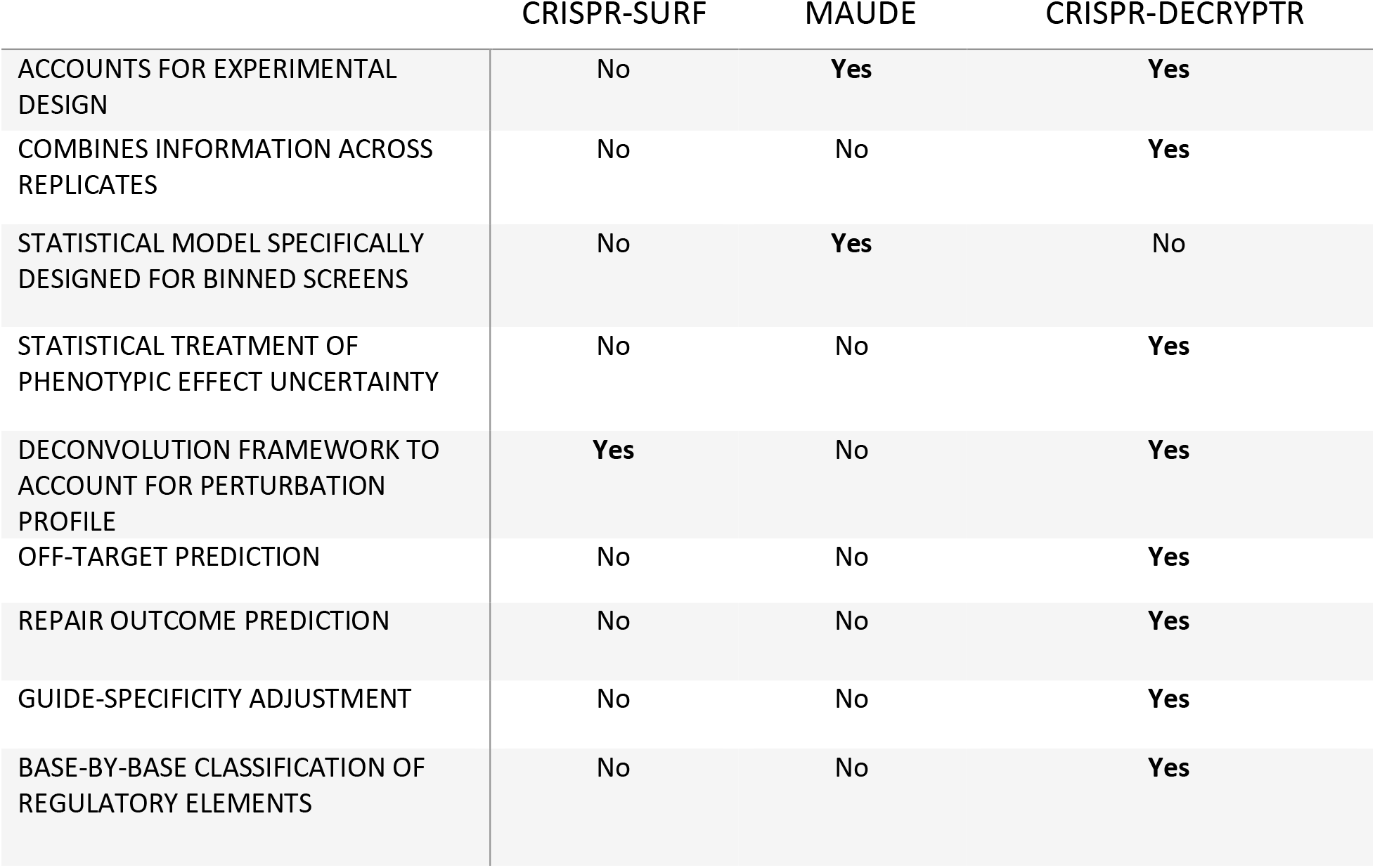
Table comparing CRISPR-SURF, MAUDE, and CRISPR-Decryptr on various attributes.

### 4.2: Discussion of Future Research

As each step in the CRISPR-Decryptr method has been designed as a modular component, they are readily adaptable as new research and technologies in gene editing emerge. Here we present our thoughts on some opportunities for future research:

#### Future research may support new generative model formulations that differ from the GLM implemented in *section 2.2*

In *section 2.2* we choose to implement a generalized linear model with a multinomial distribution due to the compositional nature of gRNA count data and simplicity of the canonical GLM framework. It is possible that different formulations could be proposed through future research and experience with the analysis of CRISPR Screens. The *infer* command’s GLM model, implemented primarily in STAN, can be readily modified.

#### Research could help with more accurate construction of convolution matrices

In this paper, we use two published works (Allen et al. and Hsu et al.) in our construction of the convolution matrix from *section 2.3*^3, 4^. It is likely there are other current or future publications that can help in more accurately predicting the perturbations of gRNAs from sequence or other features. By simply replacing the *predict* step with another means of convolution matrix construction, the CRISPR-Decryptr method is readily adaptable.

#### Research has the potential to build upon the GP deconvolution step

We believe the GP deconvolution is a powerful framework for deconvolving guide-specific effects. Future research could explore alternative kernels to the squared exponential loss that may better capture the underlying process dynamics. Additionally, if computationally feasable, the Gaussian Process parameters could be fit in a Bayesian manner.

#### Classification of regulatory elements should leverage multiple technologies

We believe it is optimal to combine results from multiple technologies in the inference of regulatory element locations. Within the scope of the paper, we compare regulatory element state calls with other genomic signals, such as histone marks and ATAC-seq. However, in practice it would be ideal to leverage information from multiple signals in one inference procedure. This could be done by adding emissions for these technologies to the HsMM in *section 2.5*.

#### The techniques presented in CRISPR-Decryptr may be of use in other analyses

The core model components of CRISPR-Decryptr also have applications to other problems in computational biology. The canonical GLM model can be extended to readouts following other distributions from the exponential family, a broad group of probability distributions that offer both discrete (binomial, poisson) and continuous (normal, exponential, gamma) supports. Additionally, the formulation of the linear predictor can be modified to account for new experimental designs or insights and account for *a priori* knowledge about model parameters. These factors establish Bayesian GLMs as a robust approach for inferring effects of genomic perturbations induced by CRISPR-Cas9 cleavage or future editing technologies. Outside of gene editing technologies, the Gaussian Process model can serve to deconvolve a variety of sparse genomic signals if the model assumptions hold on the data in question. Additionally, the HsMMs implemented in *section 2.5* can be used to decode latent state variables from a variety of genomic signals, including those with missing observations. We have implemented similar HsMM models to annotate accessible chromatin using ATAC-seq and DNAse-seq data and intend to continue the development of these models.^6^

#### Deep learning algorithms could learn generative models

Generative Bayesian models are readily interpretable and provide a statistical framework to utilize prior knowledge of uncertain systems. It is our belief that these models are invaluable for unsupervised learning tasks in computational biology, however, their proclivity to become computationally expensive can make their application to analyses of entire genomes difficult. We believe learning generative models with deep neural networks may allow for extremely fast applications of complicated models. This could be done by applying generative models to simulated or real data and using the results produced to train deep neural networks.

## APPENDIX

## A.1. Simulation Studies

## A.1.1 Log_2_ Fold-Change on Simulated Data

The Log_2_ Fold-Change is used the quantify the change in gRNA counts across conditions in some methods for noncoding and knockout screens (see section 9.1). This could be viewed as analogous to determining ‘regulatory effect’ of a targeted perturbation, as discussed further in *section 2.2* and *section 9.1*. Fold change calculations are commonly implemented in genomics and other sciences. However, the drawbacks of fold changes are well known when applied to other technologies such as RNA-seq for differential expression. The calculation is undefined with zero in the denominator and exhibits bias in that it emphasizes small denominators. Conversely, in cases where two counts are high, the ratio between them may understate the difference in counts. We demonstrate this visually in **figure 4.1** below. When considering gRNA counts, we do not believe the ratio between two measurements is an appropriate way to quantify the impact of targeted perturbations. The bias of the fold change transformation has a greater impact on correct identification of enriched guides at smaller read depths and signal strengths. We utilize forward simulations from the CRISPR-Decryptr generative model and evaluate performance of the Log_2_ fold-change to demonstrate this phenomenon. We perform 7,000 forward simulations of 10,000 gRNA counts in two conditions from the CRISPR-Decryptr at three read depths (1, 2, and 4 million reads) and calculate the Log_2_ fold change as follows:

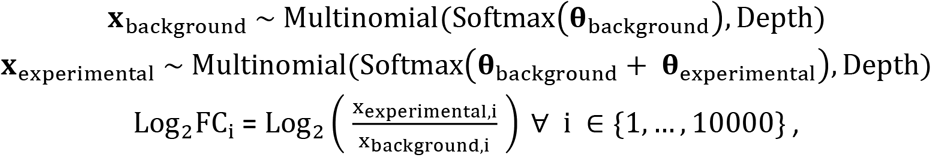

Where **x**_background_, **x**_experimental_, **θ**_background_, and **θ**_experimental_ ∈ ℝ^10000^ and Depth ∈ {1000000, 2000000, 4000000}. We set **θ**_background_ to 1, and allow 500 entries in **θ**_experimental_ to be equal to *μ* ∈ {0, 0.05, 0.10, … ‥ 1}, while the other 9,500 entries are set to zero. We consider the non-zero entries to be effects of “enriched” guides, while the zero entries are effects of “control” guides. For each read depth and value of *μ*, we calculate the expected number of simulated control guides that are within the top 95^th^ percentile of Log_2_FC, while simultaneously have background counts (x_background,i_) in the bottom 5^th^ percentile. This metric allows us to quantitatively demonstrate the tendency of the Log_2_ fold-change to appear extreme in cases where background counts are very small, as well as examine how this phenomenon is impacted by read depth and how strong the enrichment signal is in the data. In **figure 4.2**, we see that smaller read depths (Depth) and signal (*μ*) lead to higher numbers of small background counts leading to extreme Log_2_ fold-changes in simulated data.

**Figure 4.2:**
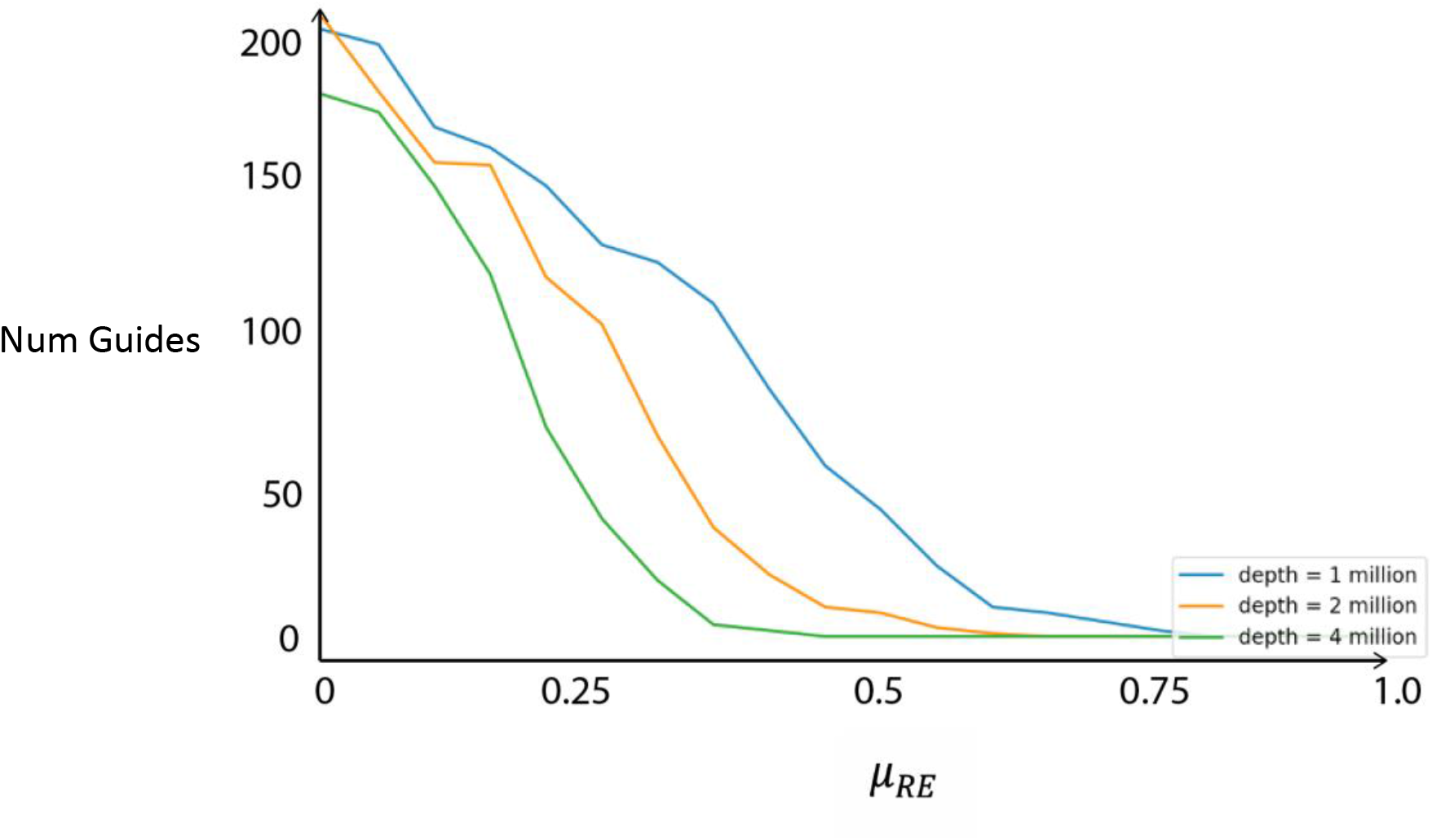
Mean number of simulated control guides that are within the top 95^th^ percentile of Log_2_FC, while simultaneously have background counts x_background,i_ in the bottom 5^th^ percentile vs. signal strength (*μ*_*RE*_) for three read depths (1 million, 2 million, and 4 million reads).

## A.1.2. Classification of Simulated Regulatory Elements

In this section, we will examine the performance of hidden semi-Markov models vs. sliding windows for classification of enriched regulatory elements. Our goal is to demonstrate the robust ability of HsMMs to fit and classify data and contrast this with the limitations of sliding-window methods with arbitrary parameters. An analysis of HsMM performance on data generated from the CRISPR-Decryptr generative model (as in the previous sections) would have been a task in classifying data using an identical model to that which it was generated from. As such, we do not generate data with a HMM or Negative Binomial state durations, but instead simulated 3kb stretches of chromatin, randomly placing up to three regulatory elements of uniform size *d*_RE_ ~ Uniform(0, 200). Bases within regulatory elements emit a normally distributed regulatory effect *e*_RE_ ~ N(*μ*_*RE*_, 1), while bases outside of regulatory elements emit a normally distributed regulatory effect *e*_RE_ ~ N(0, 1). We classify regulatory elements using the HsMM model implemented in CRISPR-Decryptr, as well as a sliding window method which calculates a Z-score for each window via Stouffer’s Method using four different parameter choices for window and step size. Iterating through 20 equally spaced *μ*_RE_ ∈ {0, 0.05, 0.1, … ,1}, we perform 500 simulations at each *μ*_*RE*_ and compare AUPR values for each method.

**Supplementary Figure 5:**
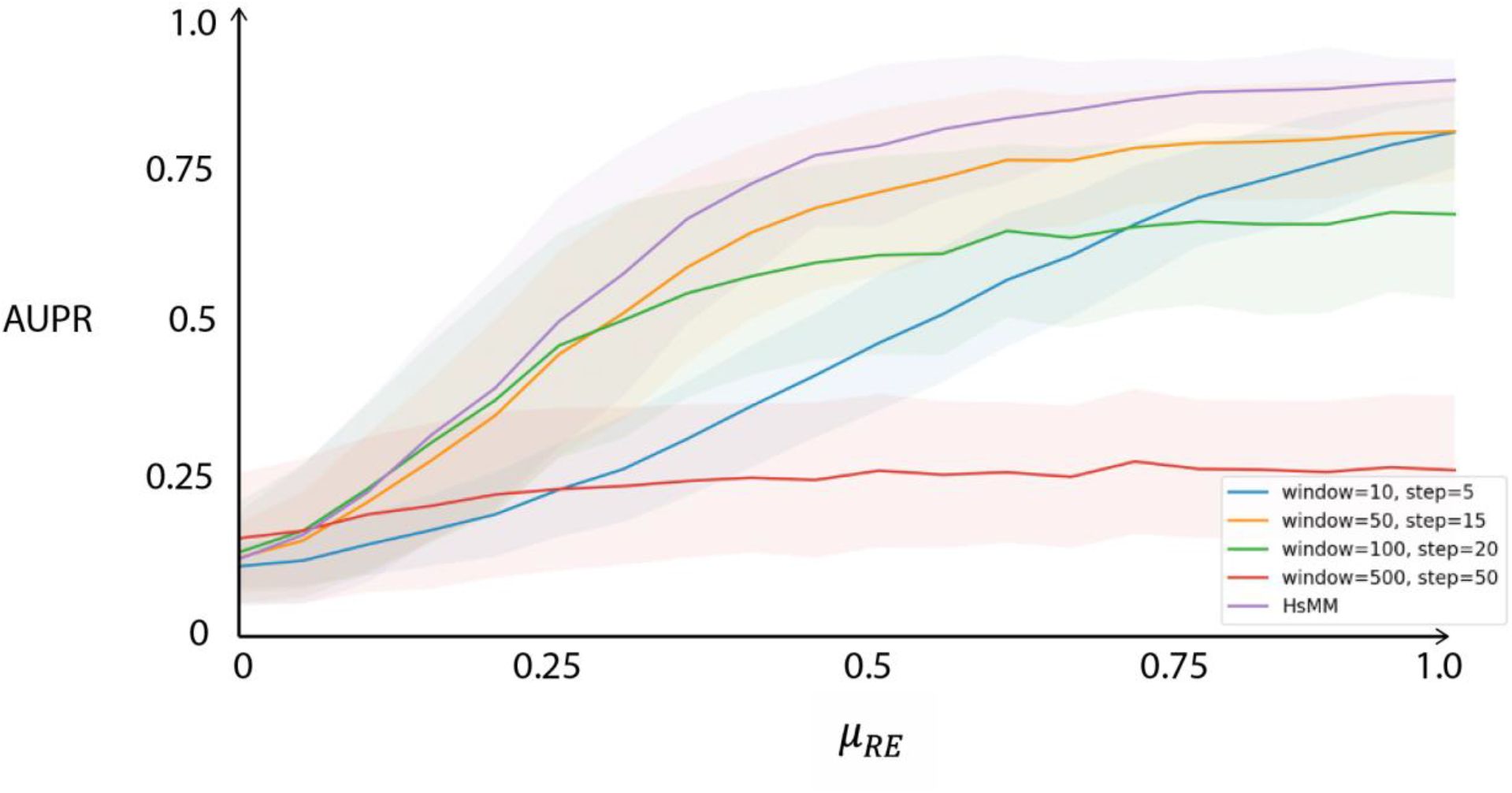
Performance of HsMM vs. Windowing Method on Simulated Data. Average Precision Recall curves across varying *μ*_*RE*_ values, the mean of the regulatory elements effect distribution. Lines indicate the mean AUPR, while the shaded region represents +/- one standard deviation.

The hidden semi-Markov model demonstrates outperformance when compared to sliding window methods, quantified by average precision recall across signal strengths. Our simulation demonstrates that the arbitrary parameters, window size and step size, greatly impact classification performance. As window size becomes large in comparison to regulatory element size (best illustrated in the **window=500, step=50** series) classification performance becomes limited even given high signal strength. At the other extreme, as window size becomes too small (best illustrated by the **window=10, step=5** series) model performance is dramatically reduced at lower signal strengths, as the model is unable to incorporate spatial dependencies of bases outside the window. Even a sliding window with the same size as the expectation of regulatory element sizes (100bps) underperforms the HsMM model. With little a priori knowledge about the size of regulatory elements, parameters for sliding window methods are generally selected arbitrarily. The HsMM model implemented in CRISPR-Decryptr does not have user-defined parameters, but instead fits latent parameters using the expectation maximization algorithm.

## A.2. Mathematical Notes

## Gaussian process based deconvolution

A continuous-time stochastic process 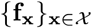 is a Gaussian process if and only if for every finite set of **x**_1_, **x**_2_, … , **x**_*M*_ in the index set of 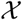

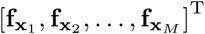

is a multivariate Gaussian random variable.

Let us consider a Gaussian process 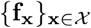 and a finite index set *X* = {*x*_*i*_ ∈ ℝ|*i* = 1, 2, … , *M*}, then

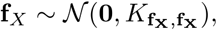

where f_*X*_ ∈ ℝ^*M*^. Moreover, let us assume that the covariance between the random variables 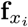 and 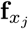 can be written as a function of the indices *x*_*i*_ and *x*_*j*_ as follows

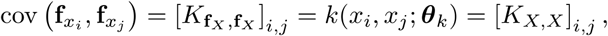

where 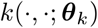 is a covariance function parameterized by the parameters in ***θ***_*k*_. Here, we use the squared exponential (SE) covariance function

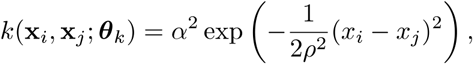

where α^2^ and *ρ*^2^ are the signal variance and the squared length-scale, respectively.

Next, let us consider the following linear transformation of f_*X*_

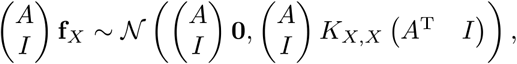

where *A* ∈ ℝ^*N* × *M*^. The resulting distribution is

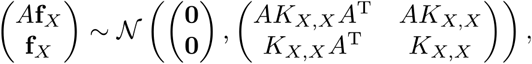

which is a joint distribution of *A*f_*x*_ and f_*x*_. Note that *A*f_*x*_ ∈ ℝ^*N*^ and *AK*_*X,X*_ *A*^T^ ∈ ℝ^*N × N*^.

Our convolution model is **m** = *A***f** + ϵ, where 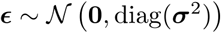. Therefore, we can write the joint dist ribut ion of **m** and **f**_*X*_ as

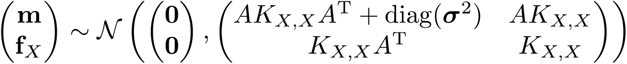

Conditioning the joint distribution on the observations **m** and their uncertainties **σ**^2^ leads to

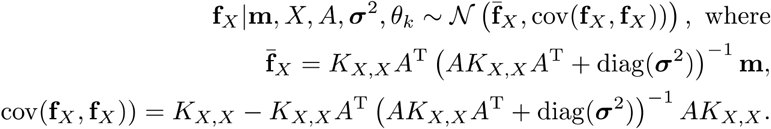

The marginal likelihood *p*(**m**|*X*, *A,* **σ**^2^, ***θ***_*k*_) is

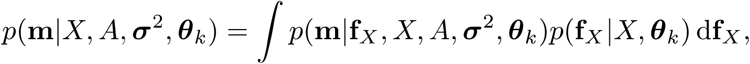

and the resulting log marginal likelihood is

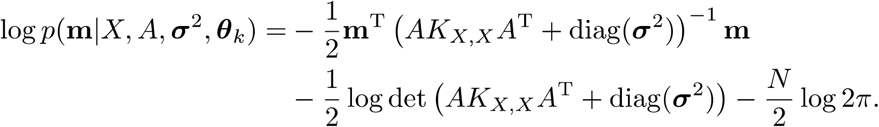

Using the log marginal likelihood, we can formulate the type-II maximum likelihood estimation of ***θ***_*k*_ as follows

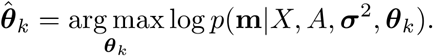

The gradient of the log marginal likelihood with respect to the parameters *α* and *ρ* is

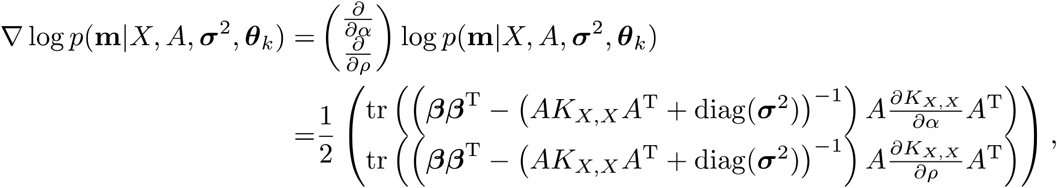

where *β* = (*AK*_*X,X*_*A*^T^ + diag(σ^2^))^−1^ **m**.

## Collapsed multinomial

Let 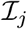, *j* = 1, 2, *N*_sets_ such that 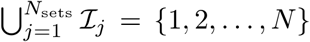 and 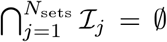. Moreover, let 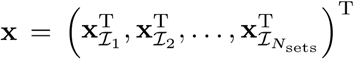 and 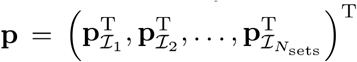. Let us consider collapsed versions of **x** and **p** as defined as follows

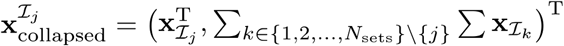

and

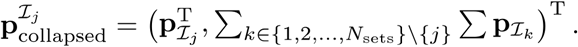

That is, both 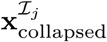 and 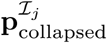 have 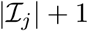 elements.

If **x** follows a multinomial dist ribution with parameters *M* and **p**

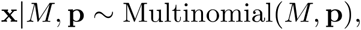

then 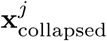 follows the multinomial distribution with parameters *M* and 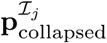

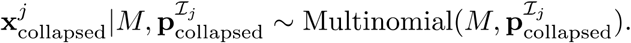

## A.3. Performance of CNN model for Repair Prediction

To demonstrate the predictive performance of the convolutional neural network used in the repair outcome prediction step, we benchmarked its performance on held-out data from the original dataset of repair outcomes from Allen et al. To do this, we randomly choose 80% of repair outcome profiles as training data, on which we trained the CNN, a 5-Nearest Neighbor Algorithm, and a predictor that guesses the average profile from the training set independent of sequence. Using remaining 20% of the dataset, we tested these three methods by predicting and evaluating their accuracy using two performance metrics: the sum of the mean squared error (MSE) and KL-divergence between predicted and realized profiles. As reported below (***Table A.3***), the CNN outperforms the other prediction methods as measured by both metrics. It is worth noting that the CNN was trained using an MSE loss function.

**Table A.3.** Sum of Mean Squared Error and KL-Divergence of realized and predicted repair profiles for CNN, Average Profile, and 5-NN algorithms. The CNN outperforms both alternative prediction methods by both metrics.

|  | MEAN SQUARED ERROR | KL-DIVERGENCE |
| --- | --- | --- |
| CONVOLUTIONAL NEURAL NETWORK | 8.83e-05 | 7.81 Bits |
| AVERAGE PROFILE | 1.23e-04 | 9.24 Bits |
| 5-NN PREDICTIONS | 1.50e-04 | 11.46 Bits |

**Figure A.3.**
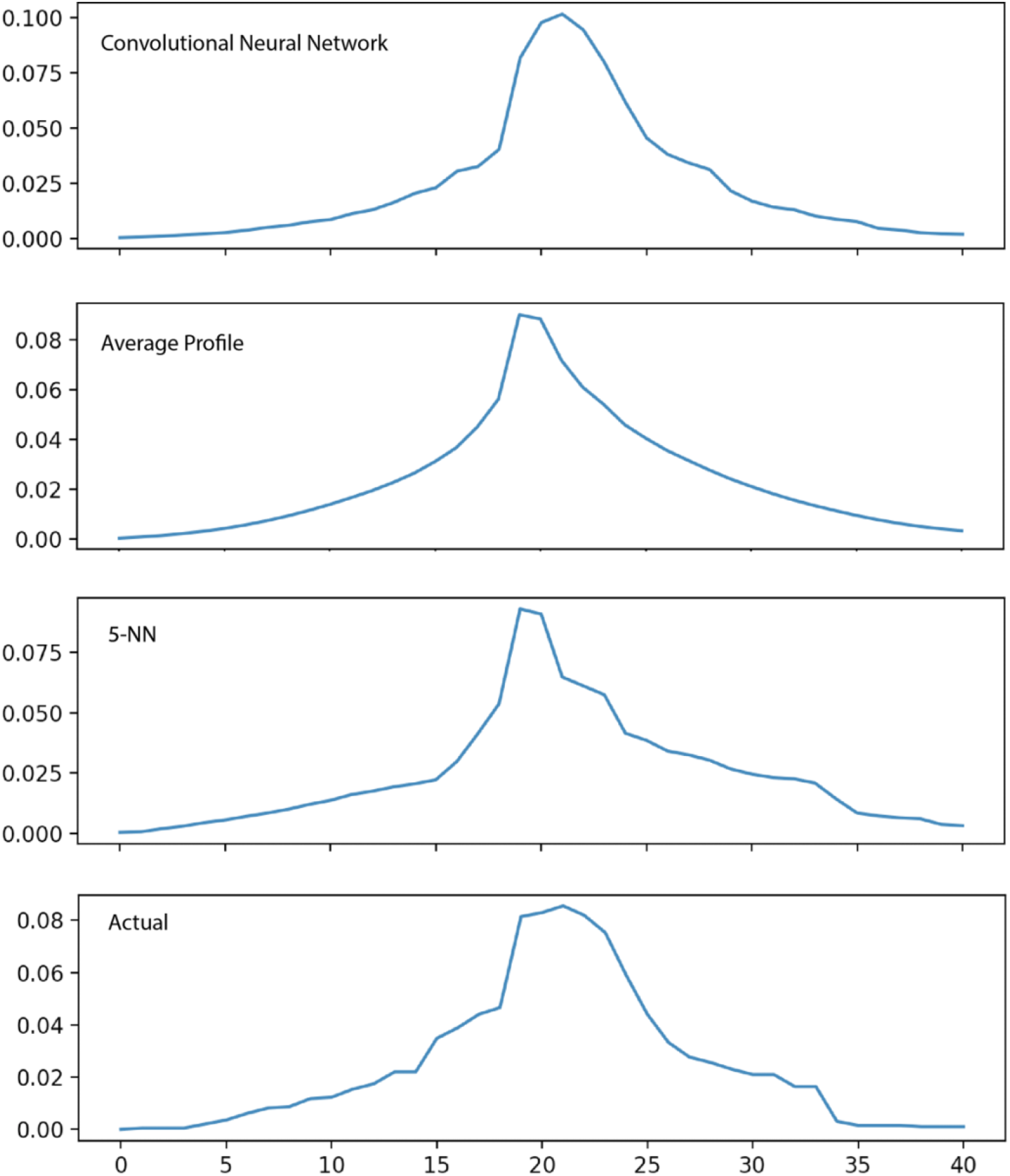
Predicted profiles and realized repair profile for the target sequence CCAGACAACAAAGCTGCCCTCGGGTAAGGATGTAGGGAGGG from the CNN model, average profile prediction, and the 5-NN.

## A.4. Notes on HsMM Hyperparameters

By default, CRISPR-Decryptr initializes a two-state HsMM with hyperparameters based on the length of the region being analyzed and the observed effect signal. Allow *N*_*bases*_ to be the length of the region and *σ*_*e*_ to be the standard deviation of the observed effect.

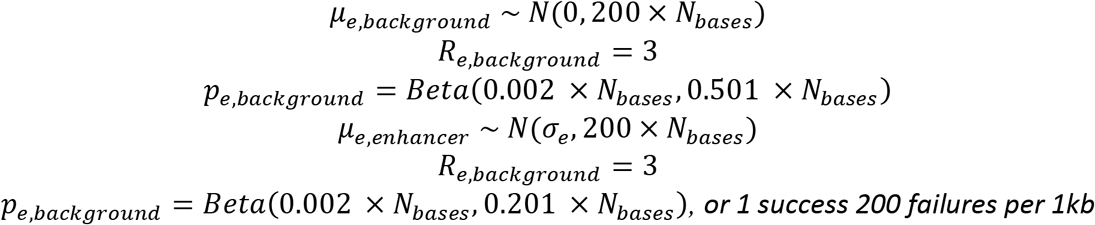

Expressed in terms of pseudocounts, hyperparameters for the parameter *p*_*e*,*s*_ represent 1 success and 500 failures per 1kb for the background state and 1 success and 200 failures per 1kb for the enhancer state.

CRISPR-Decryptr provides an argument to specify prior hyperparameters for the emission and duration distributions of the model. Unlike the default HsMM initialization, these hyperparameters are not functions of the region length or observed signal variance. Recall that the HsMM is comprised of an arbitrary number of states with of negative binomial duration, with normal emission distributions **(Figure A.4)**. The parameter r of the negative binomial can be specified by the user if desired, with a default of three. The precision of the normal distribution emission is determined at each base (*Section 2.5*). As such, parameter posteriors that will be calculated by the EM procedure are the emission means *μ*_*e*,*s*_, negative binomial parameters *p*_*e*,*s*_.

**Figure A.4.**
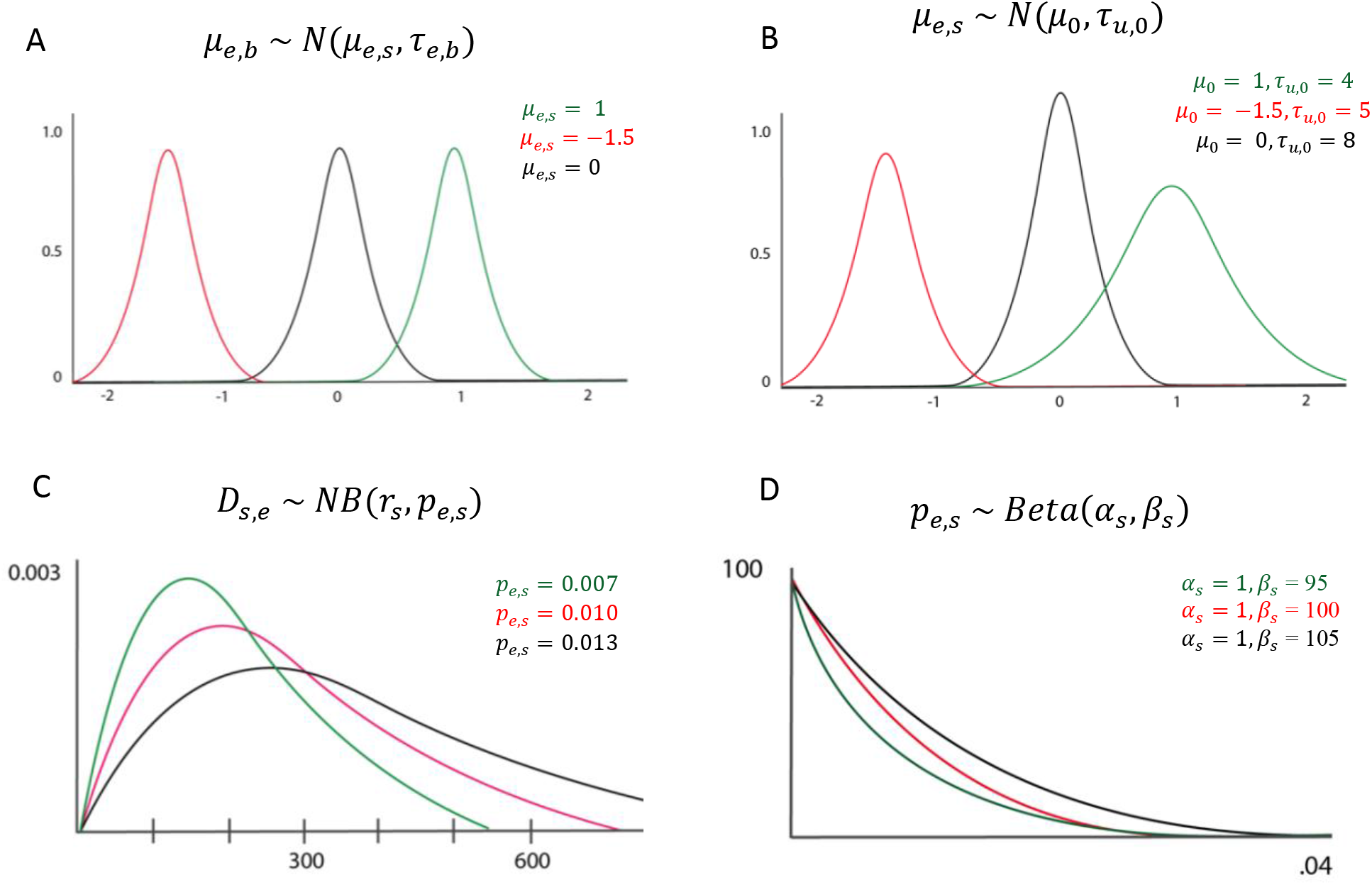
Example distributions for model parameters and priors with different priors/hyperparameters. **A:** Distribution of effect specific mean at base *b μ*_*e*,*b*_ follows a normal distribution with fixed precision at each base. **B:** Prior distribution of *μ*_*s*,*e*_ is normally distributed with hyperparameters *μ*_0_, *τ*_*u*,0_ which can be specified by the user. **C:** State duration is Negative Binomially distributed with user defined parameter *r*_*s*_ and inferred parameter *p*_*e*,*s*_. **D:** Prior distribution for parameter *p*_*e*,*s*_ follows a beta distribution.

The optional *--priors* argument takes a tab delimited file following the format of **Table A.4**. The columns of the table are defined as follows:

| Column Name: | Description: |
| --- | --- |
| *mu_0* | mean hyperparameter $\mu_0$ (figure A.4.B) of prior distribution $\mu_{s,e}$. |
| *tau_0* | precision hyperparameter $\tau_{u,0}$ (figure A.4.B) of prior distribution $\mu_{s,e}$. |
| *NB_r* | r parameter for the negative binomial duration distribution (figure A.4.C) |
| *NB_succ* | Think of NB distribution (with parameter r) as the distribution of number of succeed/fail coin flips performed (where success probability is $p_{e,s}$) before reaching r successes. This parameter represents the pseudocount of number of successes “pseudo-observed” which constructs the prior. From figure A.4.D $\alpha_s = NB_{succ,s}$. The expected state duration will be $\frac{NB_{fail} + NB_{succ}}{NB_{succ}} \times r$. |
| *NB_fail* | This parameter represents the pseudocount of number of failures “pseudo-observed” which constructs the prior (see above). |

**Table A.4.** Example of .tsv file containing hyperparameter information for a three-state model.

| state | e_mu_0 | e_tau_0 | NB_r | NB_succ | NB_fail |
| --- | --- | --- | --- | --- | --- |
| background | 0 | 5000000 | 3 | 1000 | 500000 |
| silencer | 1 | 5000000 | 3 | 1000 | 200000 |
| enhancer | -1 | 5000000 | 3 | 1000 | 200000 |

## A.5. Brief note on gRNA specificity

In the course of constructing CRISPR-Decryptr, we have made note of guide depletion when targeting certain repeat elements or regions of low sequence specificity. To our knowledge, this phenomenon has not been significantly researched, but was noted in the Canver et al. paper where the authors targeted an Alu SINE repeat in BCL11a DHS +62 (**Figure 3.1.1**). CRISPR-Decryptr accounts for guide-specificity by default, however, in our analysis of the CRISPRa CD69 screen (section 3.2), we also see gRNA depletion at repeat elements with low specificity if we disable all guide-specificity options. In this paper, we are unable to provide a biological argument as to why this would be the case in both CRISPR mutagenesis and CRISPR activation screens. However, we are confident that this phenomenon has great potential to lead to false positives. In *section 4.1* we note the propensity of the MAUDE algorithm to pick up on this “non-specificity effect,” and subsequently erroneously classify repeat regions as putative regulators. Future research appears indicated if this phenomenon is to be modeled in future methods.

**Figure A.4:**
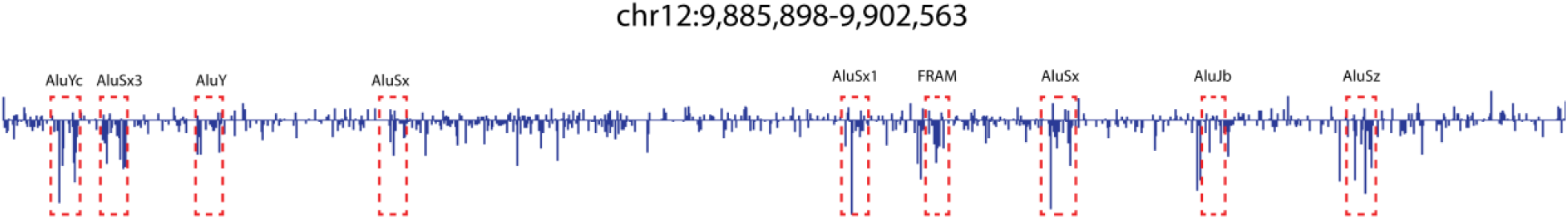
Selected region downstream of CD69 showcasing gRNA depletion at repeat elements (red boxes) as seen in the inferred regulatory effect (low bin effect, see section 3.2).

## Code Availability

CRISPR-Decryptr code and readme are located at: https://github.com/anders-w-rasmussen/crispr_decryptr

## Data Availability

gRNA count data and input files for the screens analyzed can be found at the following link:

## Acknowledgements

This research was made possible by the Simons Foundation. RB and MG acknowledge support from the following sources: NIH R01DK103358, Simons Foundation, NSF-IOS-1546218, R35GM122515, NSF CBET-1728858, and NIH R01AI130945

## Contributions

A.R. conceived of the model with guidance and oversight from R.B. and T.Ä. The inference step of the method and Gaussian Process deconvolution were formulated and coded by T.Ä. Off-target, repair outcome prediction, and the Gaussian Process / HsMM iterative procedure were formulated by A.R. The majority of HsMM code is adapted from previous work by done by M.I.G and A.R. in developing the ChromA algorithm. Rules for HsMM parameter updates and variational methods were developed by M.I.G. N.C. was instrumental in advising on high-performance computing considerations. N.S. and J.S. provided important insight into noncoding screens from the viewpoint of experimentalists and were central in bringing the need for this statistical method to A.R. and R.B.’s attention. A.R. did analyses of published data, wrote the paper supplement, and wrote the CRISPR-Decryptr software. All authors contributed to the writing of the manuscript.

## Reference Genome Data

CRISPR-Decryptr utilizes the following resources for the hg19 reference genome.

(hg19, GRCh37 Genome Reference Consortium Human Reference 37 (GCA_000001405.1)) http://hgdownload.cse.ucsc.edu/goldenPath/hg19/chromosomes/

Encode Mapability Tracks^12^: http://hgdownload.soe.ucsc.edu/goldenPath/hg19/encodeDCC/wgEncodeMapability/

